# Src activates retrograde membrane traffic through phosphorylation of GBF1

**DOI:** 10.1101/2020.08.02.233353

**Authors:** Joanne Chia, Shyi-Chyi Wang, Sheena Wee, David James Gill, Felicia Tay, Srinivasaraghavan Kannan, Chandra S Verma, Jayantha Gunaratne, Frederic A. Bard

## Abstract

The Src tyrosine kinase controls cancer-critical protein glycosylation through Golgi to ER relocation of GALNTs enzymes. How Src induces this trafficking event is unknown. Golgi to ER transport depends on the GTP Exchange factor (GEF) GBF1 and small GTPase Arf1. Here we show that Src induces the formation of tubular transport carriers containing GALNTs through promotion of a GBF1-Arf1 complex. The complex is initiated by phosphorylation on GBF1 on 10 tyrosine residues; two of them, Y876 and Y898 are located near the C-terminus of the Sec7 GEF domain. Their phosphorylation promotes partial melting of the Sec7 domain, favoring binding to the GTPase. Perturbation of these rearrangements prevent GALNTs relocation. In sum, Src promotes GALNTs relocation by favoring binding of GBF1 to Arf1. Regulation of a GEF-Arf axis by tyrosine phosphorylation appears to be a highly conserved and wide-spread mechanism.

## Introduction

Eukaryotic cells constantly regulate membrane trafficking between compartments to adjust their physiology. Thus, signaling pathways impinge on trafficking pathways at many different levels and with various outcomes. For instance, the Src tyrosine kinase has been shown to regulate Golgi membranes, in part to adjust trafficking rates in response to change in cargo load (Pulvirenti *et al*., 2008). Changes in Src activity have major effects on the morphology of the Golgi apparatus, with either contraction after Src depletion or fragmentation upon Src hyper-activation (Bard *et al*., 2003; Weller *et al*., 2010).

A specific role of the Src kinase at the Golgi is to regulate protein O-glycosylation. Src activation induces the relocation of polypeptide GalNAc transferases (GALNTs) from the Golgi to the Endoplasmic Reticulum (ER) (Bard and Chia, 2016). GALNTs initiate GalNAc type O-glycosylation and their relocation leads to a marked increase in glycosylation, which can be measured with the levels of the Tn glycan (a single GalNac residue). Increase in Tn can be detected by lectins such as HPL and VVL (Gill *et al*., 2010; Gill, Clausen and Bard, 2011). The Tn levels increase corresponds to multiple cell surface and ER resident proteins becoming hyper-glycosylated; such as for instance MMP14, PDIA4 and Calnexin (Nguyen *et al*., 2017; Ros *et al*., 2020). In short, GALNTs relocation upregulates GalNac type O-glycosylation of many proteins.

The <u>GALN</u>Ts <u>A</u>ctivation (GALA) pathway is strongly activated in breast, lung and liver cancers and presumably in most high Tn expressing tumors. GALA markedly promotes tumor growth and metastasis (Gill *et al*., 2013; Nguyen *et al*., 2017; Ros *et al*., 2020). In addition to Src, the pathway can be stimulated by the cell surface receptors EGFR or PDGFR and is controlled by a complex signaling network, including a constitutive negative regulation by the kinase ERK8 in some cell types (Chia *et al*., 2014). Src has long been implicated in tumorigenesis and tumor invasiveness and GALA is likely an important mediator of Src oncogenic effects (Chia, Tay and Bard, 2019). It is unclear how Src stimulates the transport of GALNTs from the Golgi to the ER apart that the process involves the Arf1 small GTPase and can be blocked by a dominant negative form of Arf1 (Gill *et al*., 2010).

Arf1 is part of a family of small GTPases involved in many aspects of intracellular membranes (D’Souza-Schorey and Chavrier, 2006). Arfs function in conjunction with larger proteins called GTP Exchange Factors (GEF), that mediate the transfer from GDP to GTP-bound form of the small GTPase. GEFs bind to the GDP bound form of Arfs and not the GTP bound form. All GEFs have in common a Sec7 domain (Sec7d), that specifically mediates displacement of GDP from Arf-GDP and loading with GTP. Thus, Arfs function like molecular timers, oscillating between GDP and GTP-bound forms and binding different partners (Cherfils, 2014).

There are seven subfamilies of Arf GEFs in eukaryotes, whereby two subfamilies operate at the Golgi: BIG1/2 and GBF1 (Cox *et al*., 2004). While the BIGs primarily function at the trans Golgi network and endosomal compartments, GBF1 functions at the early cis-Golgi and ER-Golgi intermediate compartment (ERGIC) and regulates Golgi to ER retrograde traffic (K. Kawamoto *et al*., 2002; Zhao, Lasell and Melançon, 2002; Zhao *et al*., 2006).

GBF1 functions mostly with Arf1. GBF1 contains five other conserved domains, two in N-terminal of the Sec7d (DCB and HUS) and three in C-terminal (HDS1 to 3). These domains are thought to mediate GBF1 recruitment to membranes, and/or regulate membrane transport (Richardson, McDonold and Fromme, 2012). Work by Melançon’s group has identified membrane-bound Arf-GDP as a factor regulating GBF1 recruitment to cis-Golgi membranes (Quilty *et al*., 2014, 2018). In addition, they proposed that the C-terminal domains HDS1 and 2 are required to bind to an unidentified Golgi receptor (Quilty *et al*., 2018). More recently, HDS1 has been shown to bind to phosphoinositides such as PIP3, PI4P and PI(4,5)P2 (Meissner *et al*., 2018).

Here, we report the set-up of an inducible Src activation system to rapidly and reliably activate the relocation of GALNTs to the ER. Acute activation of Src stimulates the formation of tubular-shaped, GALNTs-containing transport carriers and results in increased O-glycosylation levels. Src activation induces the recruitment of GBF1 at the Golgi, increased binding of GBF1 to Arf1 and a transient up-regulation of Arf1-GTP levels. By mass spectrometry, we found Src directly phosphorylates GBF1 on ten residues, including residues Y876 and Y898, located within and close to the C-terminus of the Sec7d respectively. In silico modelling and directed mutagenesis suggest important conformational changes that promote binding to Arf1. As supported by additional mutants, Y876 phosphorylation induces partial melting of an alpha-helix in Sec7d, increasing binding affinity to Arf1. Y898 phosphorylation appears to release an interaction between Sec7d and the linker domain in C-terminal. We propose that tyrosine phosphorylation increases GBF1 affinity for Arf-GDP on Golgi membranes and promotes the relocation of GALNTs through tubular transport intermediates.

## Results

### Src8A7F chemical activation induces rapid GALNTs relocation to the ER

ER relocation of GALNTs can be monitored with ease by measuring total Tn levels using staining with *Helix pomatia* lectin (HPL) (Hammarström *et al*., 1977; Gill *et al*., 2010) (Figure 1A). High levels of GALA are found in a majority of samples from malignant tumors. By contrast and for unknown reasons, most cancer cell lines in vitro show limited levels of GALA (Gill *et al*., 2013). A more marked relocation can be induced in cell lines by transfecting a plasmid expressing an active form of Src . However, this approach implies an uncontrolled increase of Src activity over several hours and tends to result in fragmentation of the Golgi apparatus (Bard *et al*., 2003).

**Figure 1:**
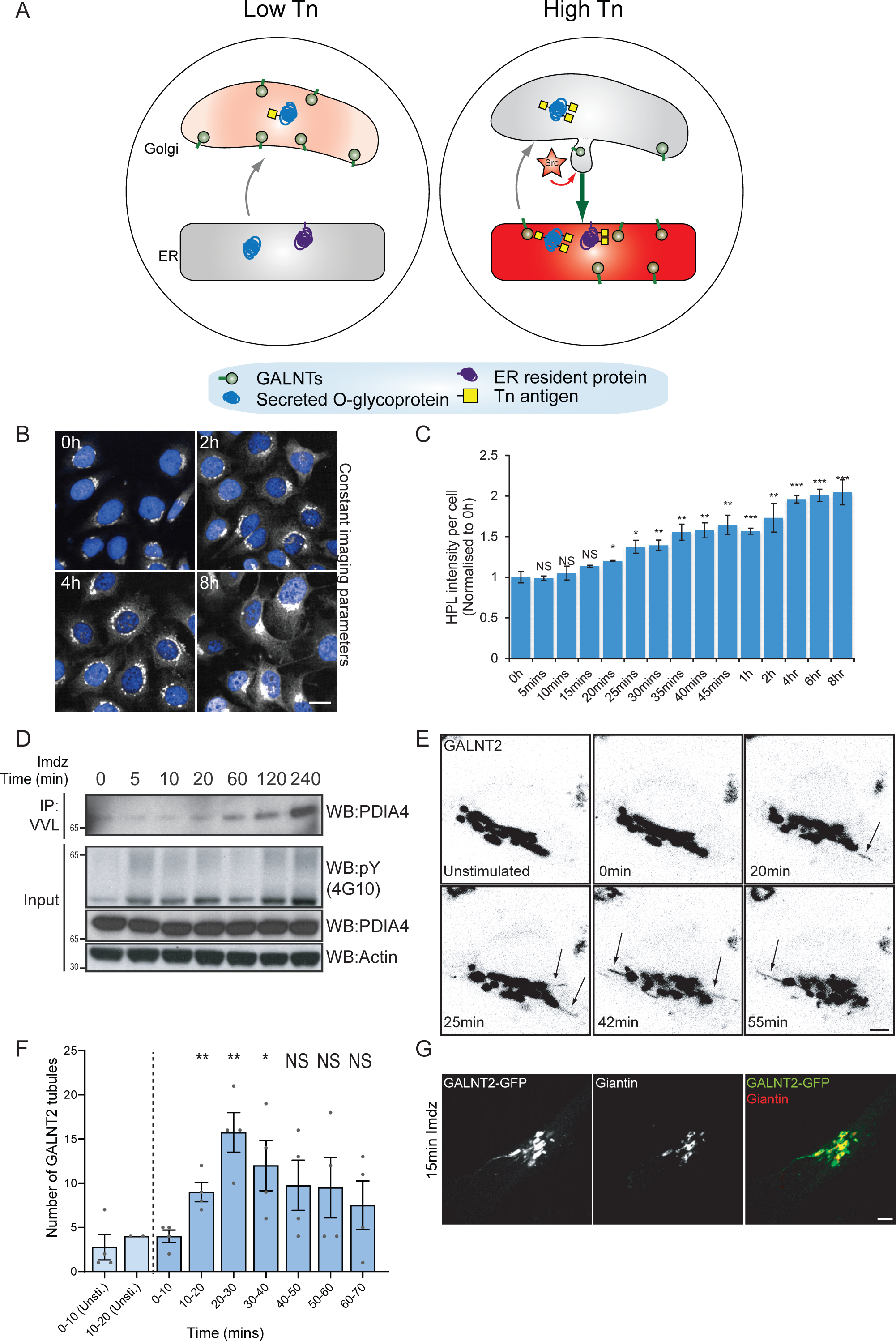
Src8A7F chemical activation induces GALNTs relocation to the ER in tubular carriers. (A) Schematic of the GALA pathway, the red coloring represents the anti-Tn lectin staining. (B) Representative images of HPL staining of Tn in Hela-IS cells after 5 mM imidazole (imdz) stimulation. Scale bar: 20 μm. (C) HPL staining intensity per cell normalized to untreated control cells (0h). Three replicate wells per experiment were quantified. (D) Representative immunoblot analysis of VVL immunoprecipitation of cell lysate after 5 mM imdz treatment of HEK-IS cells. (E) Still images of time-course analysis of GALNT2-GFP expressing Hela-IS cells stimulated with 5 mM imdz. Scale bar: 5 μm. (F) Quantification of the number of GALNT2 tubules emanating from the Golgi over various time pre-imdz (light blue bars) and post-imdz treatment (dark blue bars). Tubules were counted manually over 10 minute windows in four independent cells. (G) Fixed GALNT2-expressing Hela-IS cells were stained for the Golgin Giantin. Values on graphs indicate the mean ± SD. Statistical significance (p) were measured by two-tailed paired t-test.*, p < 0.05, **, p<0.01 and ***, p < 0.001 relative to untreated cells.

To obtain a better control of the Src-induced GALNT relocation, we sought to rapidly activate Src. The Cole group has demonstrated that a mutant form of Src, Src(R388A,Y527F) or Src8A7F for short is a mostly inactive kinase that can be chemically rescued and activated by imidazole (Qiao *et al*., 2006) (Figure S1A). After generating a stable Hela cell line expressing this Inducible <u>Sr</u>c, HeLa-IS, we verified that imidazole treatment induces an increase in total tyrosine phosphorylation. By comparison with the overexpression of constitutively active point mutant SrcE378G (SrcEG), the increase in tyrosine phosphorylation remained moderate (Figure S1B).

In terms of GALA, imidazole treatment induced a two-fold increase in total Tn levels after two hours with a pattern of Tn staining a mix of Golgi and ER, suggesting a measured GALNT relocation (Figure 1B-C, S1C). Staining pattern of the Golgi marker Giantin was not affected, indicating that the Golgi organisation was not overly perturbed. Tn increase was relatively modest compared to the expression of SrcEG (Figure S1D) or ERK8 depletion in HeLa cells (Chia *et al*., 2014). Imidazole treatment in wild type HeLa cells had no effect on Tn (Figure S1E). We also observed that the effects of Src8A7F activation was reversible: imidazole washout after 24 hours of treatment resulted in significant reduction of Tn levels within 1 hour (Figure S1F-G).

To evaluate the physiological relevance of the inducible Src activation system, we compared it to a stimulation with platelet-derived growth factor (PDGF). PDGF binding to the PDGF receptor usually results in Src activation (Thomas and Brugge, 1997). Stimulation with 50 ng/ml PDGF yielded around two-fold increase in total Tn levels (Figure S1H-I), similar to that observed with imidazole treatment. Hence, Src8A7F rescue recapitulates the levels of response to growth factor stimulation. The advantage of imidazole rescue is a reliable Src activation, whereas the effect of PDGF stimulation tends to be influenced by cell culture conditions (Chia, Tay and Bard, 2019). Similar data was obtained in another cell line HEK293 that stably expresses Src8A7F (HEK-IS). HEK-IS recapitulated the results obtained with HeLa-IS, providing an alternative model for biochemical experiments.

We further verified whether increased Tn levels were due to relocation of GALNTs to the ER. Direct observation of GALNTs in the ER is technically challenging because of the dilution and dispersion factor involved (Chia, Tay and Bard, 2019). However, GALNTs presence in the ER results in O-glycosylation of ER resident proteins such as PDIA4, which is more readily quantified (Nguyen *et al*., 2017; Chia, Tay and Bard, 2019). We measured the effects of Src8A7F imidazole rescue on the glycosylation of PDIA4, using *vicia villosa* lectin (VVL) immunoprecipitation followed by PDIA4 blotting and quantified a five-fold increase of PDIA4 glycosylation after four hours (Figure 1D). Overall, the Src8A7F system provides a measured activation of the GALA pathway, without breakdown of the Golgi structure and with tight kinetic control.

### Acute activation of Src induces GALNTs-containing tubules at the Golgi

To visualise GALNTs relocation, Hela-IS cells stably expressing GFP-GALNT2 were imaged by time-lapse microscopy after imidazole stimulation. In unstimulated conditions, GALNT2 was mostly confined at the Golgi. Upon imidazole addition, GFP-positive tubules started to emanate from the Golgi as soon as 10 mins after stimulation, their numbers reaching peak around 20-30 min then decreasing to slightly above unstimulated conditions (Figure 1E-F, movie 1). The tubules often detached and moved away from the Golgi, suggesting effective transport (Figure 1F).

This phenomenon was also observed after PDGF stimulation where GALNT2 tubules emerged from the Golgi after ∼15 mins (Figure S1J, movie 2). In some particularly responsive cells, the tubules were forming at a high rate and eventually led to a marked reduction of GALNT2 levels in the Golgi. Of note, similar tubules were also observed upon drug inhibition of ERK8, a negative inhibitor of GALNTs relocation (Chia *et al*., 2014). The tubules were deprived of the peripheral Golgi protein Giantin (Figure 1G). In addition, they appeared deprived of the chimeric Golgi enzyme beta 1,4-galactosyltransferase (GALT) tagged with mCherry (Figure S1K).

### Src activation does not increase COPI recruitment on Golgi membranes

We next wondered whether tubules formation was dependent on the COPI coat. We first measured if COPI was recruited at the Golgi upon Src activation using staining for the beta-subunit of COPI. Surprisingly, using both high resolution microscopy as well as quantitative automated high-throughput confocal microscopy, we observed no increase but instead a mild reduction of COPI intensity at the Golgi between 5 to 20 minutes after Src activation (Figure S1L-N). These results suggest that COPI is not playing a driving role in the formation of tubules at the Golgi, consistent with previous reports about the formation of retrograde-directed tubular intermediates at the Golgi (Bottanelli *et al*., 2017). Since COPI vesicles formation cannot be readily observed, our observations suggest that GALNTs retrograde traffic to the ER is mediated instead by tubules emanating from the Golgi and seceding into transport carriers.

### Transient Src activation increases Arf1-GTP levels

Tubular carriers involved in retrograde traffic have been described previously and recently shown to depend on the small GTPase Arf1 (Bottanelli *et al*., 2017). We previously reported the requirement of Arf1 for GALNTs relocation (Gill *et al*., 2010; Chia *et al*., 2014). By contrast, the small GTPase Arf3 is thought to act primarily in anterograde traffic at the TGN (Sztul *et al*., 2019). Consistently, siRNA knockdown of Arf1 resulted in significant reduction of Tn levels upon imidazole treatment and Arf3 knockdown had little effect (Figure 2A-B, S2E).

**Figure 2:**
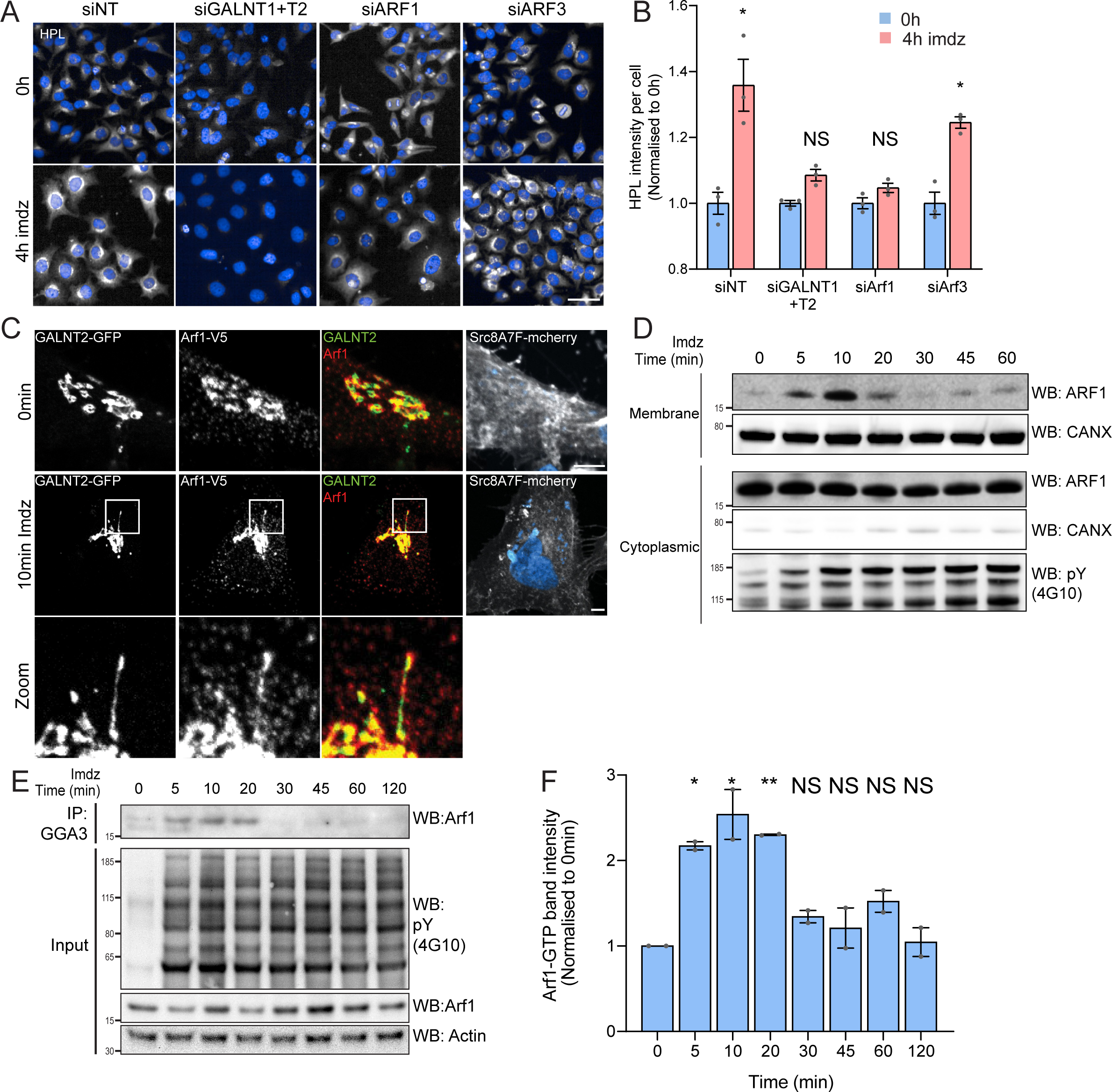
Src activation stimulates GTP loading and membrane recruitment of Arf1. (A) Representative images of HPL staining in Hela-IS treated with various siRNA before and after 4 hours of imdz treatment. siNT refers to non-targeting siRNA and siGALNT1+T2 refers to co-transfection of GALNT1 and GALNT2 siRNAs. Images were acquired under constant acquisition settings. Scale bar: 50 μm. (B) Quantification of HPL staining intensity per cell normalized to the respective untreated cells (0h) for each siRNA treatment. Three replicate wells per experiment were measured. (C) Representative images of GALNT2-expressing Hela-IS cells stained for Arf1 before and after 10 minutes of stimulation with 5 mM imdz. Images were acquired at 100x magnification. Scale bar: 5 μm. (D) SDS-PAGE analysis of cytoplasmic and membrane levels of Arf1 after imdz stimulation. CANX refers to blotting for ER resident Calnexin. The blots were generated with the same exposure and repeated twice. (E) SDS-PAGE analysis of GTP loaded Arf1 after pulldown with GGA3 beads after imdz treatment in HEK-IS cells. (F) Quantification of Arf1-GTP levels in (E). Two experimental replicates were measured and values were normalised to untreated cells (0h). Values on graphs indicate the mean ± SD. Statistical significance (p) were measured by two-tailed paired t-test.*, p < 0.05, **, p<0.01 and ***, p < 0.001 relative to untreated cells. NS, non significant.

Arf1 has been involved in the formation of retrograde tubular carriers at the Golgi (Beck *et al*., 2008; Krauss *et al*., 2008; Bottanelli *et al*., 2017). We wondered if Arf1 was present on GALNT2 tubules, however antibody staining was too faint to be conclusive. We thus generated Hela-IS stably co-expressing GALNT2-GFP and C-terminal V5 tagged Arf1 (Arf1-V5). The small V5 tag was selected to minimize functional interference and we picked a clone that expresses moderate levels of Arf1-V5 (Jian *et al*., 2010). We found that in unstimulated cells, Arf1-V5 localises both at the Golgi and in peripheral cytosol (Figure 2C). Upon Src8A7F activation, Arf1-V5 appeared to be recruited at the Golgi and localised on the GALNT2 tubules, almost throughout the structure (Figure 2C, S2A). To confirm the membrane recruitment of Arf1, we isolated cytosolic and membrane proteins to measure the levels of membrane-bound Arf1 and found Arf1 membrane-association increased within 5 minutes, peaked at 10 minutes and began to fall after 20 minutes while the cytosolic pool remained relatively constant (Figure 2D,S2D). The results so far suggested that Arf1 is activated by Src, suggesting an effect on Arf1-GTP levels. We measured them using pull-down with the binding domain of the Arf1 effector GGA1 in HEK-IS (Dell’Angelica *et al*., 2000; Yoon, Bonifacino and Randazzo, 2005). Strikingly, Arf1-GTP levels were increased more than two-fold within 5 minutes of imidazole induction (Figure 2E-F). Interestingly, Arf1-GTP levels subsided after 30 minutes of stimulation despite continuous Src activity. Surprisingly, transient expression of SrcEG for 18 hours resulted in a marked decrease in the amount of Arf1-GTP (Figure S2B-C).

Altogether, the data indicates that Src activation at the Golgi results in a transient increase in GTP loaded Arf1 and recruitment at the Golgi. Given the reported increased affinity of Arf-GTP for membranes, the switch to GTP-bound form might explain the increase in membrane-bound Arf (Pasqualato, Renault and Cherfils, 2002; Nawrotek, Zeghouf and Cherfils, 2016).

### GBF1 is required for Arf-GTP formation, GALNT relocation and tubules formation

GTP loading of Arf1 at the Golgi is regulated by GBF1 (Kazumasa Kawamoto *et al*., 2002; Zhao *et al*., 2006). We previously reported that lowering GBF1 levels reduces GALNTs relocation in cells where GALA has been induced by ERK8 depletion(Chia *et al*., 2014). Following Src activation in imidazole-treated HEK-IS, GBF1 RNAi mediated knockdown resulted similarly in a significant reduction of Tn levels (Figure S3A-B) and PDIA4 glycosylation (Figure S3C). These results indicate that GBF1 is mediating the burst of GALNTs relocation induced by Src8A7F and suggest that GBF1 may also control Arf-GTP burst.

To test this, we reasoned that increased GBF1 expression together with Src activation should enhance Arf-GTP formation. Expression of a GFP-tagged form of GBF1 (GFP-GBF1) alone enhanced Arf1-GTP levels in HEK-IS (Figure 3A). Strikingly, upon imidazole stimulation, GTP loading was further increased by nearly 3 fold within 10 minutes of induction (Figure 3A-B). The effect was transient and Arf1-GTP returned to pre-stimulation within 45 minutes, indicating similar dynamics as with wild-type levels of GBF1.

**Figure 3:**
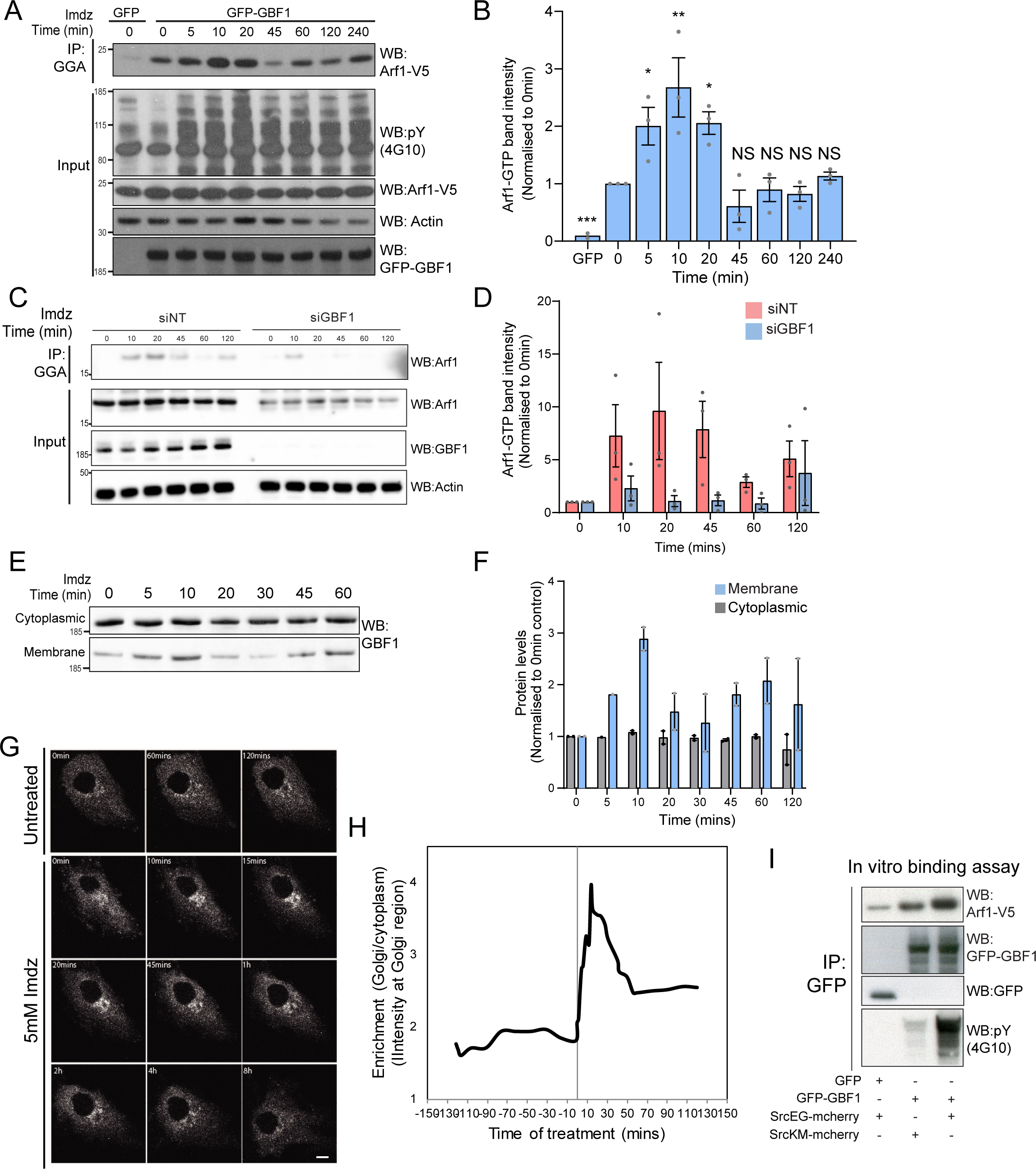
Src activates the ARF-GEF GBF1. (A) Representative SDS-PAGE analysis of Arf1-GTP levels in HEK-IS cells expressing GFP or GFP-GBF1. GGA pulldown was performed as in Figure 2E. (B) Quantification of Arf1-GTP levels in three independent experiments. (C) SDS-PAGE analysis of cytoplasmic and membrane levels of GBF1 after imdz stimulation. (D) Quantification of two independent experiments shown in (C). Values presented were normalised to untreated cells (0h). (E) Still images of the time-lapse movie of GBF1-GFP in Hela-IS cells stimulated with 5 mM imdz. Scale bar: 10 μm. (F) Quantification of the ratio of Golgi to total cytoplasmic levels of GBF1 before and after imdz treatment in time-lapse shown in (E). (G) SDS-PAGE analysis of the levels of Arf1-V5 bound to GFP or GFP-GBF1 IP from cells expressing inactive SrcKM or active SrcEG in an *in vitro* binding assay. Two experimental replicates were tested and quantified in Figure S3E.

We next tested whether GBF1 depletion would block Arf-GTP production upon Src activation. This experiment is delicate as GBF1 depletion tends to affect cell adhesion and Golgi organisation, so we timed it carefully to when GBF1 levels start to drop (Figure 3C). In these conditions, imidazole stimulation resulted in a reduced Arf-GTP formation in GBF1 depleted cells (Figure 3D).

We next tested whether Src also stimulates GBF1 recruitment to membranes in HEK-IS. Strikingly, the GBF1 membrane pool increased by roughly 3 fold within 5-10 minutes, while total GBF1 remained constant. This increase was relatively transient, decreasing after 10 minutes of imidazole treatment (Figure 3E-F). Using time-lapse microscopy of GFP-GBF1 in Hela-IS, we observed GBF1 recruitment at the Golgi complex. In sync with the biochemical experiment, GBF1 Golgi levels increased rapidly, peaking at ∼ 10 minutes, and decreasing after ∼20 minutes (Figure 3G-H).

GBF1 recruitment at the Golgi depends on binding to membrane bound Arf-GDP (Quilty *et al*., 2014). Thus, we wondered whether phosphorylation by Src might increase GBF1 affinity for Arf1. To test this idea, we isolated Arf1-V5 from a cell lysate and added, in the presence of GDP, GFP-GBF1 immunoprecipitated on beads from cells expressing also active or inactive Src (Figure S3D). After 1h incubation, beads were washed and the amount of Arf1 bound to beads quantified by immunoblotting (Figure 3I). By comparison with inactive SrcKM, SrcEG induced a 2 fold increase in binding (Figure S3E). Given this net increase, we next tested binding of immunoprecipitated GFP-GBF1 with bacterially produced and purified Arf1-del17-His in the presence of GDP. Arf1-del17-His, a recombinant protein deleted of the first 17 amino-acids, is able to bind GDP in the absence of phospholipids (Kahn *et al*., 1992). As with Arf1-V5, phosphorylated GBF1 displayed increased binding to purified Arf1-del17-His (Figure S3F).

We next imaged GALNT2 tubules formation in GBF1-depleted cells. After imidazole stimulation, control cells displayed tubules formation, sometimes very abundant, while it was drastically reduced in GBF1 depleted cells (Figure S3G-H).

Altogether, our results indicate that Src activation activates induces an increase of affinity between GBF1 and Arf1-GDP, resulting in a transient but marked recruitment of GBF1 at the Golgi, GALNT2 tubules formation and subsequently a wave of Arf1-GTP.

### GBF1 protein is phosphorylated by Src on at least 10 tyrosine residues

We next tested directly whether Src phosphorylates GBF1. After imidazole stimulation of HEK-IS cells, GBF1 was immunoprecipitated and probed with an antibody specific for tyrosine phosphorylation, revealing an increase within five minutes that was sustained for two hours (Figure 4A). As the signals both for GBF1 and its phosphorylation were not very marked, we also tested GFP-GBF1 expressing HEK-IS cells; obtaining similar results (Figure 4B). We also transiently co-expressed GFP-GBF1 with SrcEG and SrcKM mutants in HEK293T cells. In such conditions, the difference in phosphorylation levels were very marked (Figure 4C). Similarly, endogenous GBF1 was phosphorylated by SrcEG in HeLa cells (Figure S4A). Finally, we tested GBF1 phosphorylation in a third cell line: NIH3T3vSrc are mouse fibroblasts that have transformed with a viral, oncogenic and constitutively active mutant of Src (Vogt, 2012). These cells display significantly higher levels of GALA than their normal counterparts. They also display four times as much phospho-GBF1 (Figure 4D). To test whether phosphorylation was direct, we immunoisolated GFP-GBF1 from HEK293 cells and added recombinant Src. In the presence of ATP, GFP-GBF1 displayed marked phosphotyrosine levels compared to controls, indicating that GBF1 is a direct substrate of Src (Figure 4E, S4C).

**Figure 4:**
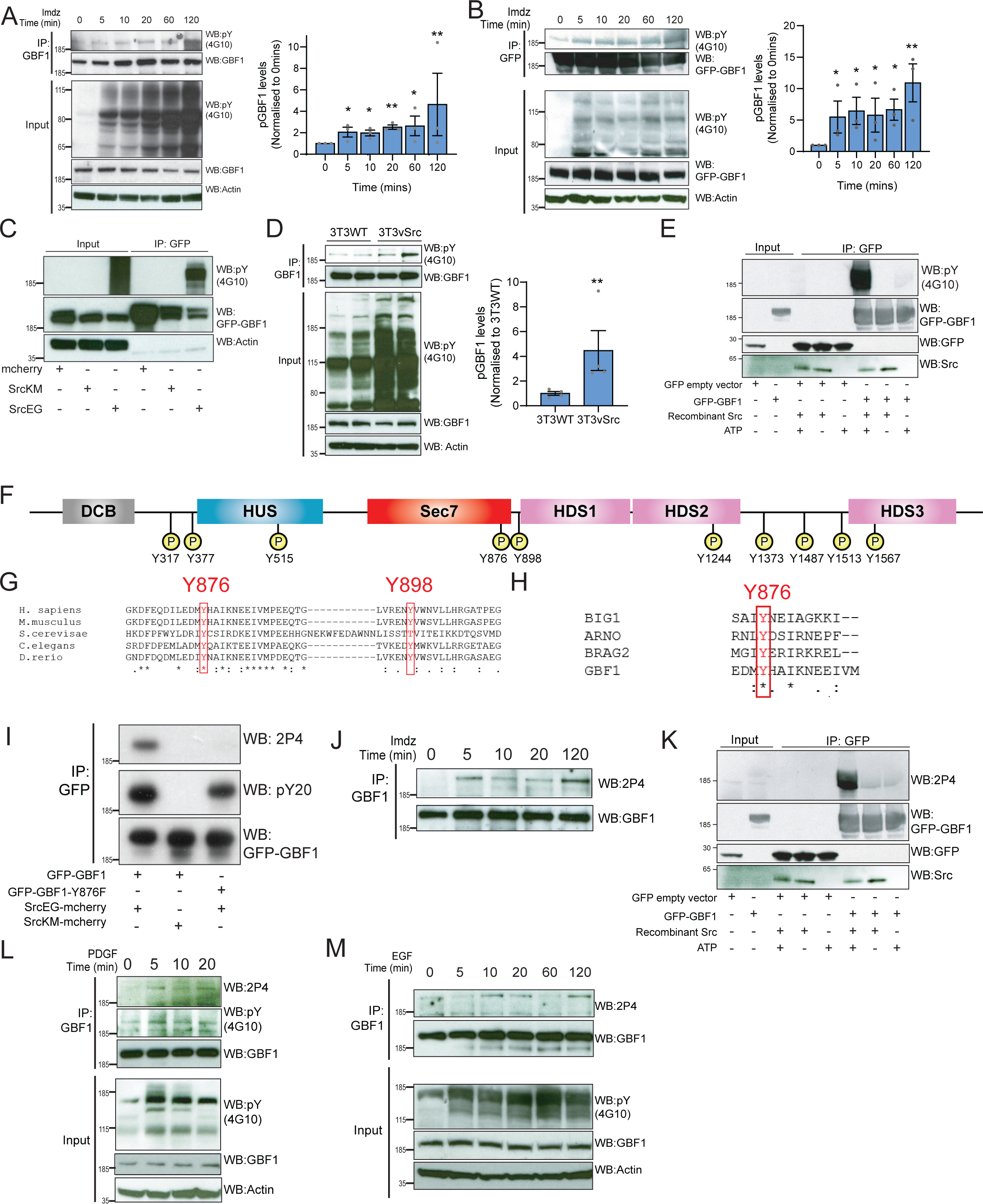
Src phosphorylates two tyrosines Y876 and Y898 at the C-terminus of the GBF1 Sec7d. (A) SDS-PAGE analysis of phospho-tyrosine (pY) levels in endogenous GBF1 using HEK-IS cells after imdz treatment. Quantification of pY-GBF1 in three replicates shown on the graph (right). (B) SDS-PAGE analysis of pY levels on GFP-GBF1 IP from HEK-IS cell line after imdz treatment. Quantification of pY-GBF1 in three replicates shown on the graph (right). (C) SDS-PAGE analysis of pY in GBF1 in HEK293T cells expressing either inactive SrcKM or active SrcEG. (D) SDS-PAGE analysis of pY levels on endogenous GBF1 IP from wild type and vSrc transformed NIH3T3 cell lines. Quantification of pY-GBF1 in four replicates from two independent experiments shown on the graph (right). (E) Quantification of pY in GBF1 after *in vitro* phosphorylation. Immunoprecipitated GFP or GFP-GBF1 was incubated with recombinant Src protein in the presence or absence of nucleotide ATP. (F) Schematic of the 10 tyrosine residues in GBF1 that were identified by targeted mass spectrometry after exposure to Src. (G) Amino acid sequence alignment of GBF1 from various species. The sequences of GBF1 at Y876 and Y898 of *H. sapiens* (NP_004184) was aligned with that of *M. musculus* (NP_849261), *S. cerevisae* (NP_010892), *C. elegans* (NP_001255140) and *D. rerio* (XP_009305378), revealing conservation of both residues. (H) Y876 is conserved and observed to be phosphorylated in other GEFs BRAG2, ARNO and BIG1 based on the Phosphositeplus database. (I) SDS-PAGE analysis of wild type GFP-GBF1 or GFP-GBF1-Y876F mutant immunoprecipitated from HEK293T cells expressing inactive SrcKM or active SrcEG. Phosphorylation at Y876 was marked by the 2P4 antibody. (J) SDS-PAGE analysis of Y876 phosphorylation on endogenous GBF1 IP from HEK-IS cell line over various durations of imidazole treatment. (K) Y876 phosphorylation of GBF1 in an *in vitro* phosphorylation assay. (L) SDS-PAGE analysis of the total and Y876 phosphorylation on endogenous GBF1 in Hela cells over the duration of 50ng/ml PDGF stimulation. (M) SDS-PAGE analysis of Y876 phosphorylation on endogenous GBF1 in A431 cells over time of 100ng/ml EGF stimulation. Values on graphs indicate the mean ± SD. Statistical significance (p) measured by two-tailed paired t-test.*, p < 0.05, **, p<0.01 and ***, p < 0.001 relative to untreated cells. NS, non significant.

To map the phosphorylation sites, GFP-GBF1 from HEK293 cells expressing active SrcEG was extracted from a gel separation and digested using trypsin and AspN. Phosphopeptides were analysed with tandem mass spectrometry, revealing ten phosphorylated tyrosine residues with high confidence (Figure 4F, S4D). All the phosphopeptides identified were clearly increased in the presence of SrcEG but mostly not detectable with inactive SrcKM expression (Figure S4D).

### Src phosphorylates two tyrosines at the C-terminus of GBF1 GEF domain

10 phosphosites are challenging to study in parallel. We were particularly interested in the residues Y876 and Y898 because they are located respectively within the GEF/Sec7d and on a C-terminal loop connecting Sec7d and the HDS1 domain (Figure 4F). It suggested they could be directly involved in the regulation of GBF1 GEF activity.

A database search revealed that Y876 is conserved in GBF1 homologues from all species investigated from yeast *S. cerevisae* to *H. sapiens* (Figure 4G). Y898 is conserved in GBF1 from all species considered except yeast (Figure 4G, S4E). In contrast, the other phosphorylation sites are mostly conserved in vertebrates and Y317 is only present in human GBF1 (Figure S4E). We next compared Sec7 domains of different ARFGEFs and found Y876 to be highly conserved, while Y898 appeared specific for GBF1. Interestingly, a search on PhosphositePlus database indicates that multiple GEFs, including Brefeldin-Resistant Arf-GEF 2 (BRAG2, IQSEC1), Brefeldin A-Inhibited Guanine Nucleotide-Exchange Protein 1 (BIG1) and Cytohesin 2 (ARNO) can be phosphorylated on the tyrosine analogous to Y876 (Figure 4H).

We subsequently used targeted SILAC to quantify the intensity of phosphorylation on each site in HEK293 transiently transfected (Ong *et al*., 2002). GBF1, in the presence of SrcEG, displayed about 180-fold increase in phosphorylated Y876 peptide (DFEQDILEDMyHAIK) and 100-fold in increase in phosphorylated Y898 peptide (ENyVWNVLLHR) compared to SrcKM expressing samples, suggesting these sites are the major phosphorylation targets (Figure S4F-H).

### Phosphorylation at Y876 in endogenous GBF1 is confirmed with a specific antibody

We next aimed to generate antibodies specific for phospho-Y876 and Y898. While our efforts on Y898 were unsuccessful, we obtained a monoclonal antibody named 2P4 after immunisation with a Y876-containing phosphopeptide that reacted with phospho-GBF1 (Figure S4I). To verify specificity, wild type GFP-GBF1 and mutant GFP-GBF1(Y876F) were co-expressed with SrcEG, immunoprecipitated and probed with 2P4. While wildtype GBF1 showed strong reactivity and total phosphotyrosine levels were moderately affected, the band was completely abolished in the Y876F mutant (Figure 4I).

We used 2P4 to assess the kinetics of Y876 phosphorylation after Src activation in the HEK-IS system. Similar to overall phosphotyrosine levels, phospho Y876 was detected within five minutes of imidazole treatment and persisted for two hours (Figure 4J). Similar results were obtained with GFP-GBF1 (Figure S4J). We also verified that Y876 is a direct target of Src using the in vitro phosphorylation assay (Figure 4K, S4K).

We also tested Y876 phosphorylation after growth factor stimulation. Starting with serum starved Hela cells stimulated with PDGF, endogenous GBF1 was immunoprecipitated and phospho-Y876 found to display similar kinetics to generic GBF1 tyrosine phosphorylation (Figure 4L). A431 cells, which express high levels of EGFR, were stimulated with 100ng/ml of EGF (Fernandez-Pol, 1985). Similar to PDGF with HeLa cells, phospho-Y876 was upregulated within 10-20 minutes (Figure 4M). To review, Y876 is a major site of phosphorylation by Src and is modified in physiological conditions of GALA activation.

### Phosphorylation at Y876 and Y898 regulates GBF1 binding to Arf1-GDP

To establish the functional importance of Y876 and Y898, we generated single and double Y to F mutants at position 876 and Y898 (Y876F, Y898F, Y876.898F). Hela-IS cells were transfected with wild type or mutant GBF1 and tested for Arf1 GTP loading. Similar to previous data, Arf1-GTP loading increased by ∼2.5-fold within 10 minutes of imidazole treatment with wild type GBF1 expression (Figure 5A-B). This stimulus was nearly abolished with the expression of the Y876F mutant, while residual activation was observed for the Y898F mutant. As expected the double mutant could not be stimulated and even basal Arf1-GTP levels were reduced.

**Figure 5:**
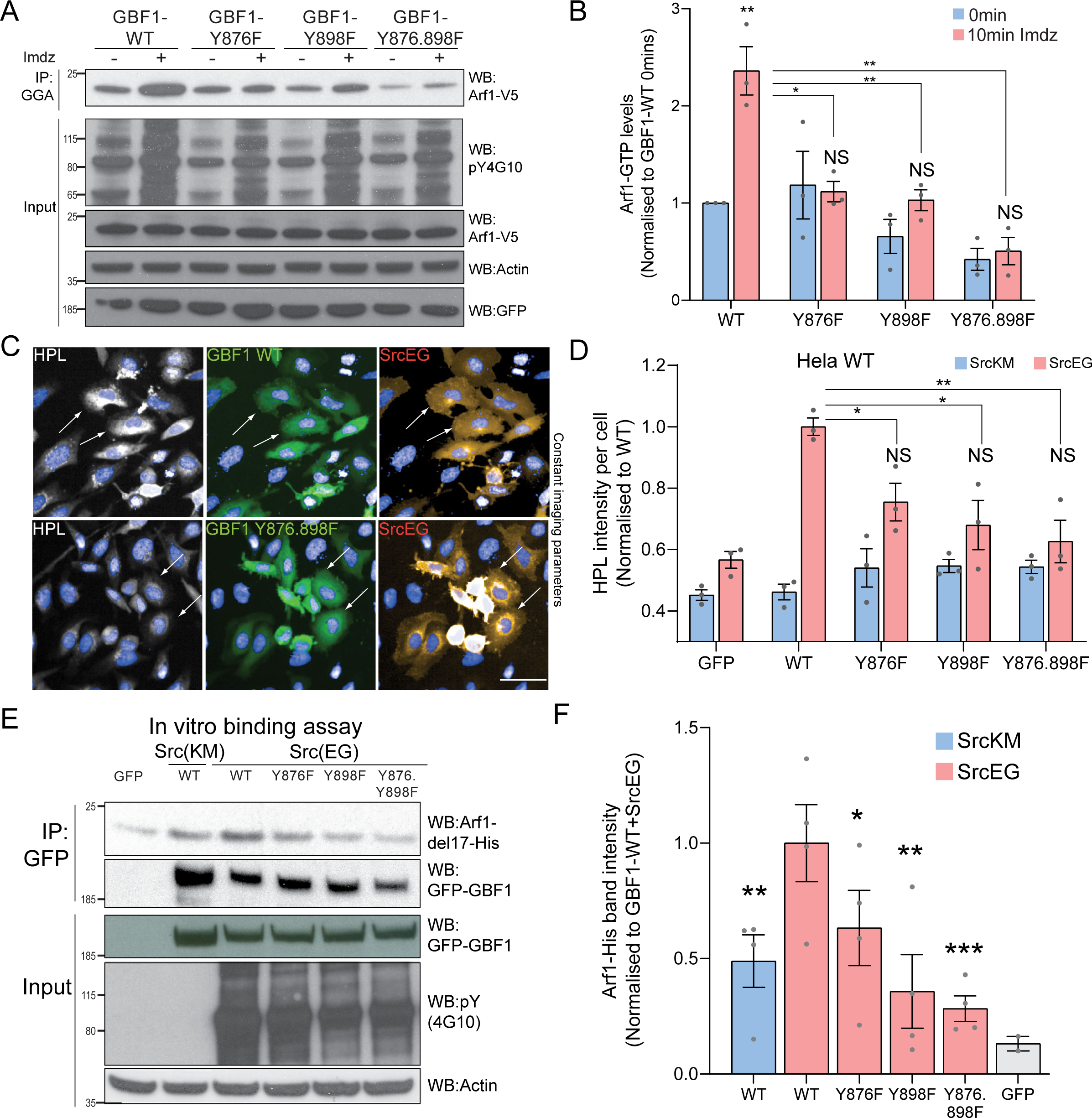
Phosphorylation at Y876 and Y898 regulate GEF activity of GBF1. A) SDS-PAGE analysis of GTP loaded Arf1 at 0min (-) and 10min (+) imidazole treatment in HEK-IS cells expressing wild type GBF1, GBF1-Y876F, GBF1-Y898F or GBF1-Y876.898F mutants. (B) Quantification of Arf1-GTP loading levels in (A) from three independent experiments. Values were normalised to untreated cells (-) expressing wild type GBF1. (C) Representative images of HPL staining in Hela cells co-expressing wild type GBF1 or GBF1-Y876.898F mutant with active SrcEG. Scale bar: 50 μm. (D) Quantification of HPL staining levels of cells co-expressing wild type or mutant GBF1 with inactive SrcKM (blue bars) or active SrcEG (pink bars). Values were from three replicates. (E) SDS-PAGE analysis of the levels of recombinant Arf1-His bound to wild type or mutant GFP-GBF1 IP from cells expressing inactive SrcKM or active SrcEG in an in vitro binding assay. (F) Quantification of the levels of bound Arf1-His. Values were from four experimental replicates and normalised to wild type GBF1 IP from cells expressing active SrcEG from each experiment. Immunoprecipitated GFP protein used as a negative control for non-specific binding with GFP (grey bar). Values on graphs indicate the mean ± SD. Statistical significance (p) measured by two-tailed paired t-test.*, p < 0.05, **, p<0.01 and ***, p < 0.001 relative to untreated cells or to 10-min imdz treated cells expressing wild type GBF1. NS, non significant.

Next, we tested the effect of Y to E phospho-mimetic mutations for both sites (Y876E and Y898E). Surprisingly, we observed a reduction of basal GTP loading levels by more than 70% (Figure S5A-B). This resonates with the effect induced by expression of the constitutively active SrcEG (Figure S2B). It suggests that the Y-to-E mutants are able to recapitulate some of the effects of Src on the Golgi.

Next, we tested whether GALNT relocation was affected: wild type or mutant GBF1 were co-expressed with active SrcEG and Tn levels were measured. GBF1 mutants GBF1-Y876F or GBF1-Y898F significantly repressed Tn levels in SrcEG expressing cells (Figure 5C and 5D).

These results suggest that these GBF1 mutants can act partially at least as dominant-negative and prevent the formation of retrograde transport carriers.

As phosphorylated GBF1 increasingly binds to Arf1 (Figure 3G, S3F), we assumed that loss of phosphorylation on Y876 and Y898 would affect Arf binding. We used the in-vitro binding assay with purified Arf1-del17 and immuno-purified GFP-GBF1 and indeed, single mutants of Y876 or Y898 had reduced Arf1 binding by 40% and 60% respectively, and by more than 70% for the double mutant (Figure 5E,F). In fact, the double mutant had lower binding affinity than wild type GBF1 expressed together with the kinase-dead SrcKM.

Altogether, these results indicate that phosphorylation on tyrosines Y876 and Y898 drives an increase of affinity of GBF1 for Arf1-GDP, in turn increasing Arf1-GTP levels and promoting GALNTs relocation.

### Phosphorylation on Y898 probably releases a Sec7d-HDS1 intramolecular interaction

We next wondered how the phosphorylations affect GBF1’s GEF activity. Y898 is located in the linker region between the Sec7d and HDS1 domains of GBF1. While the Sec7d structure of GBF1 has not been resolved, GBF1 Sec7d shares at least 65% homology with several other GEFs, so it can be modelled relatively accurately. Unfortunately, this is not the case for HDS1 for which there is no structural information. We could model the Sec7d and the linker domain, using GBF1 sequence and the resolved structures of the GEFs ARNO, Cytohesin-1 and Grp1. In this model, the linker is located close to a pocket of negatively charged residues in the Sec7d (Figure S6A). Molecular dynamics revealed a repulsion of the linker away from Sec7d after phosphorylation (Figure S6B, Movie S1). This suggests that phosphorylation could relieve an intramolecular interaction between the Sec7 domain and the HDS1 domain. Both ARNO and Grp1 contain a domain in C-term of Sec7d that interacts and inhibits the GEF activity, for ARNO by about 14-fold (Stalder *et al*., 2011) and about 7-fold for BIG (Richardson, McDonold and Fromme, 2012). While for both proteins, the C-term domain is a Pleckstrin Homology (PH), we hypothesise that the HDS1 of GBF1 could similarly inhibit the Sec7d and phosphorylation at Y898 would alleviate this Sec7d-HDS1 inhibition.

### Phosphorylation of Y876 partially unfolds GBF1 Sec7d domain, increasing affinity for Arf1

By contrast with Y898, Y876 is located within the Sec7d. There are 10 alpha helices in Sec7d and Y876 is present on Helix J (Cherfils *et al*., 1998). We modeled the interaction of GBF1 with Arf1 based on available structures and observed that Helix J is protruding in the interface between the two proteins (Movie S2). When we simulated Y876 phosphorylation, the negative charge was attracted by the positively charged residues Arginine 843 (R843) and lysine 844 (K844) at the end of Helix H. This, in turn, led to a partial unwinding of Helix H, leading to an extension of the loop between helices H and I (Figure 6A, Movie S3). This partial unfolding and loop extension would result in better bond formation between Sec7d and Arf1 (Figure 6B, S7A). This translates into a reduced free binding energy (Figure 6C, S7B) and an increased probability of buried surface area between the Sec7d and Arf1 (Figure 6D) in the phosphorylated state. Higher buried surface area indicates tighter packing interactions and thus, higher affinity between the molecules. Thus, the loop extension induced by phosphorylation is predicted to favor the interaction with Arf1, a result consistent with our in vitro binding assay results with phospho-GBF1 (Figure 3G, 5G).

**Figure 6:**
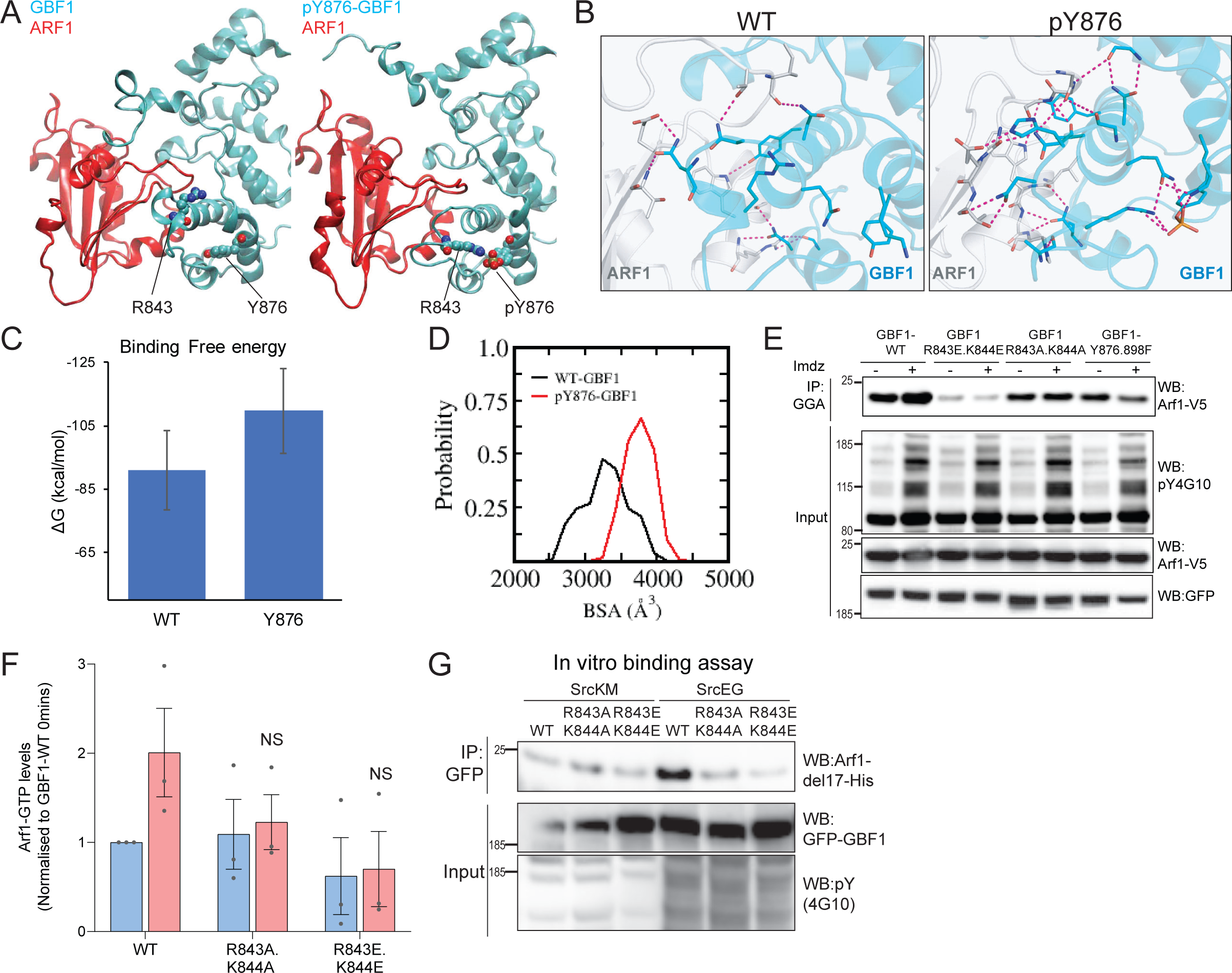
Phosphorylation on Y876 increases GBF1 Sec7d affinity for Arf1. (A) Structural basis for the binding of unphosphorylated GBF1 Sec7d and Y876 phosphorylated GBF1 Sec7d with Arf1. Cartoon representations of a representative conformation extracted from the molecular dynamics (MD) simulations of the unphosphorylated GBF1 Sec7d (left of panel A; cyan) and Y876 phosphorylated GBF1 Sec7d (right of panel A; cyan) bound to Arf1 (red colour). MD suggests the unwinding of the helix H to form an extended loop between Helix H and I through increased attractions between positive charges on R843 and K844 on the loop with the negative charges on phosphorylated Y876. The Sec7d of GBF1 (blue) in turn, interacts more with Arf1 (red). (B) Residues involved in GBF1: Arf1 inter-protein are shown as sticks, and the H-bonds highlighted as black dashed lines. The Sec7d was shown in blue while Arf1 protein in grey. Refer to Figure S7A for the identities of the residues. (C) Estimation of the free energies (ΔG) of the interactions between the unphosphorylated GBF1 Sec7d and Arf1 and between the Y876 phosphorylated GBF1 Sec7d and Arf1. Calculations carried out using the MMPBSA approximations averaged over the conformations generated from MD simulations of the complexes; larger negative values represent higher affinities. GBF1 Sec7d has a higher affinity for Arf1 when it is phosphorylated at Y876. (D) Probability distributions of the buried surface area (BSA) between GBF1 Sec7d and Arf1. (E) SDS-PAGE analysis of GTP loaded Arf1 at 0min (-) and 10min (+) imidazole treatment in HEK-IS cells expressing wild type GBF1, Y876.898F and the HI loop mutants. (F) Quantification of Arf1-GTP loading levels in (E) in three experimental replicates. Values were normalised to untreated cells (-) expressing wild type GBF1. (G) SDS-PAGE analysis of the levels of recombinant Arf1-His bound to wild type or the HI loop mutants GFP-GBF1 IP from cells expressing inactive SrcKM or active SrcEG in an *in vitro* binding assay. Values on graphs indicate the mean ± SD. Statistical significance (p) measured by two-tailed paired t-test. NS, non significant.

Since the model predicts that the positive charges on either R843 and K844 are important for the conformational change induced by phosphorylation, we mutated these sites into glutamic acids (R843E.K844E) or neutral charges with alanine (R843A.K844A). We next tested if these HI loop mutants have an impact on GTP loading on Arf1 in cells. The introduction of negative charges in the R843E.K844E mutant resulted in a massive reduction in basal cellular Arf1-GTP levels by about 90% (Figure 6E-F). On the other hand, the mutant with neutral charges (R843A.K844A) did not have an effect on the basal Arf1-GTP levels. However, when we stimulated Src8A7F with imidazole, the cells expressing the R843A.K844A mutant did not upregulate Arf1 GTP loading (Figure 6E-F). These results indicate that the positively charged residues in the loop between Helix H and I are required for the Y876 phosphorylation effect.

The model predicts that blocking the partial unfolding of helix H would reduce the interaction of GBF1 with Arf1 (Figure 6B, S7A). As expected, we found that the mutants (both E and A) were insensitive to SrcEG in terms of enhanced binding to Arf1-GDP (Figure 6G).

We next tested whether blocking helix H unfolding would prevent Src induced GALNT relocation to the ER. HPL staining intensities in cells co-expressing constitutively active SrcEG and wild type or the HI loop mutants (both A and E forms) were measured. The HI loop mutants resulted in a significant reduction in Tn levels, at levels similar to GBF1-Y876F (Figure S7C-D). A similar reduction was observed in Hela Src8A7F cells stimulated with imidazole (Figure S7E-F).

Altogether, these results strongly support a model where the phosphorylation on Y876 induces a partial melting of the Sec7d helix H, which in turn facilitates GBF1 binding to Arf1-GDP.

## Discussion

In this report, we describe how a tyrosine kinase, Src controls a key regulator of membrane trafficking, GBF1 and in turn mediates the movement of Golgi enzymes. The relocation of the GALNTs from the Golgi to the ER, its induction by the Src kinase and its critical importance for tumor growth have been established previously (Gill *et al*., 2010, 2013; Gill, Clausen and Bard, 2011; Nguyen *et al*., 2017; Ros *et al*., 2020). However it remains unclear how Src induces this relocation.

Our observations establish a critical link in the GALA pathway. They also led us to propose a new model of how GBF1 functions and is regulated during Golgi to ER traffic. Indeed, the classical model of GBF1 role in retrograde traffic is that it serves primarily to generate Arf-GTP, which in turn recruits the COPI coat and produces coated vesicles (Beck *et al*., 2009; Donaldson and Jackson, 2011; Spang, 2013). Several of our data do not fit this construct for GALNTs relocation.

First, we do not observe a surge of COPI vesicles upon Src activation. Using the Src8A7F mutant provides a system to rapidly activate GALA and a burst of relocation of GALNTs. We did not observe an increase of COPI at the Golgi, while we could clearly detect GBF1 being recruited. We also could not observe COPI vesicles, instead live imaging revealed GALNT2-filled tubules. These tubules fit well the requirement of specific GALNTs transport carriers as a reporter for the enzyme B4GAL-T1 and Giantin are excluded from them. Several groups have reported before the imaging of tubules-derived transport carriers emanating from the Golgi (Sengupta *et al*., 2015; Bottanelli *et al*., 2017). By contrast, it should be noted that the models implicating COPI vesicles are mostly based on genetic experiments in yeast and not yet supported by in vivo imaging (Cosson and Letourneur, 1994; Letourneur *et al*., 1994; Szul and Sztul, 2011).

The second point concerns Arf-GTP. We were initially expecting that Arf-GTP would drive the formation of tubules, as several studies have proposed a role for Arf-GTP in affecting membrane topology and even prompting tubules in vivo (Beck *et al*., 2008; Krauss *et al*., 2008; Bottanelli *et al*., 2017). Indeed, the burst of Arf-GTP after Src activation would tend to support this notion.

Arf depletion also clearly reduces GALNT relocation. However, the rest of the data is rather ambiguous. Mainly, there is a lack of correlation between Arf-GTP levels and GALNTs transport. Indeed, while Src activation raises transiently the levels of Arf-GTP, long term activation of Src actually depletes Arf-GTP (see Figure S2B). In these conditions, GALNTs levels in the ER are maintained, presumably by active Golgi to ER transport. Indeed, Src activity and GBF1 are still required at these time points to maintain GALNT activity in the ER.

In addition, the expression of wild-type GBF1 results in significantly increased Arf-GTP, without any observable GALNTs relocation (see Figure 3A and Figure S5A). Of course, it is possible that the pools of Arf-GTP generated in these two conditions are in different locations or with different local concentrations, but it clearly indicates that total Arf-GTP levels are not a good predictor of transport activity.

Interestingly, the first descriptions of retrograde traffic tubules were obtained after Brefeldin-A (BFA) treatment, in conditions where Arf-GTP cannot be formed (Sciaky *et al*., 1997). It was later discovered that GBF1 is a key target of BFA, which induces a stable GBF1-Arf-GDP complex on membranes, suggesting that GBF1 is directly involved in BFA-induced tubule formation but that Arf-GTP is not directly required (Niu *et al*., 2005). At this stage, the exact role of Arf-GTP in the formation of GALNTs tubules remains to be clarified. It remains of course well-established that Arf-GTP plays important roles in COPI coat recruitment and in maintaining Golgi integrity.

How can we combine the data presented into an alternative coherent model? A key insight is the increased affinity of GBF1 for Arf1-GDP as observed in the in vitro assays. This affinity increase is also supported by the modeling work and the additional GBF1 mutants based on the modeling. The improved binding can explain the surge in Arf-GTP after Src activation. Indeed, assuming that the GEF catalytic activity, the k_cat_ of the reaction, is not changed, the increased formation of GBF1/Arf-GDP complexes will mechanically lead to more Arf-GTP (Figure 7A). The increased levels of GBF1/Arf-GDP complex can also explain the increase of GBF1 at the Golgi after Src activation. In the alternative proposed model, the increased affinity of GBF1 for Arf-GDP leads to longer lasting complexes on Golgi membranes, which in turn drive the formation of tubules (Figure 7B). This mechanism would have some similarity with a BFA treatment, which stabilises GBF1 on Golgi membranes (Niu *et al*., 2005). The difference is that upon phosphorylation, GBF1 can still induce Arf nucleotide conversion. GDP to GTP conversion, in this model, would induce the disassembly of the complex and induce the release GBF1 from the membranes, limiting the formation of tubules. Admittedly, further experimentation will be required to fully test this alternative model.

**Figure 7:**
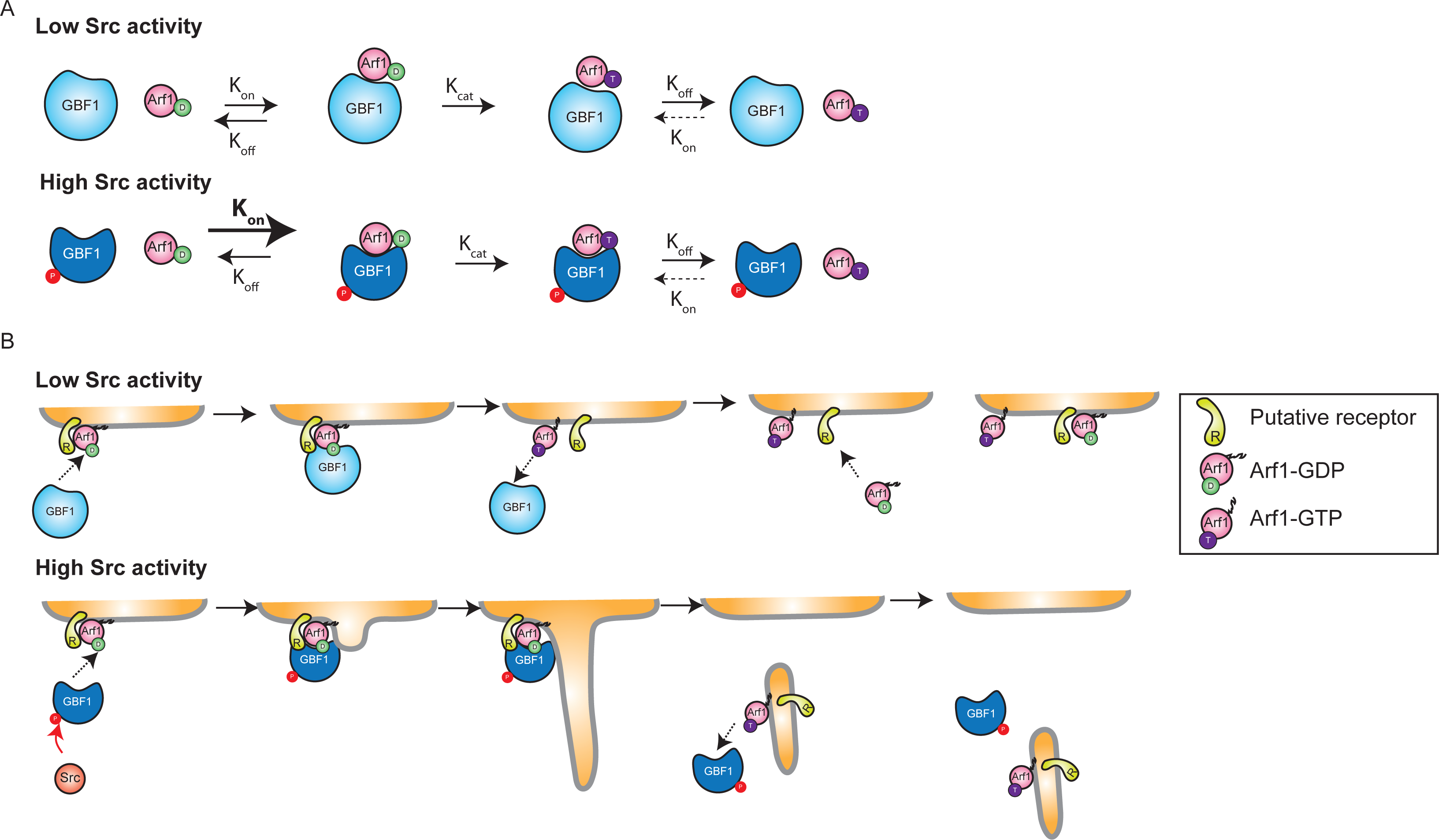
Models of GBF1 phosphorylation effects on binding to Arf1 and on the formation of tubules. (A) Increased binding affinity of phosphorylated GBF1 to Arf1-GDP promotes the formation of GBF1-Arf1-GDP complexes at the Golgi. This in turn promotes the formation Arf1-GTP. (B) Model for a self-limiting mechanism of tubules and Arf-GTP formation. (Top) In conditions of low GBF1 phosphorylation, GBF1-Arf1-GDP interaction is too transient to promote tubules formation, Arf1-GTP can accumulate in membranes, for instance when GBF1 is overexpressed. (Bottom) In conditions of high GBF1 phosphorylation, GBF1-Arf1-GDP interaction is stabilised and leads to transport carrier formation. As a result, the receptor for Arf1-GDP on Golgi membranes is progressively depleted and limits the transport of GALNTs.

It is not clear how the GBF1/Arf-GDP complex might induce tubules. It is known that their formation depends on microtubules and associated motors, so maybe GBF1 recruits specific motors (Echard *et al*., 1998). Interestingly, the GBF1/Arf1 complex has been shown to interact with the microtubule motor Miro at mitochondria(Walch *et al*., 2018).

A striking observation is that tubules show a peak of formation after Src activation, as do Arf1 and GBF1 recruitment on membranes and Arf1-GTP levels, all of which rise roughly between 10-20 minutes and decrease after 30 minutes. We also often observe a signal rebound for most of these parameters after 60-100 minutes although we have not quantified it precisely. These dynamics suggest a self-limiting process is involved. GBF1 phosphorylation itself is sustained over hours, so its affinity for Arf-GDP should be maintained. Membrane-bound Arf-GDP is probably not the limiting factor, since we observe a marked increase in total membrane-bound Arf after Src activation. This observation suggests that membrane-bound Arf-GDP is rapidly replenished from the cytosolic pool after conversion to Arf-GTP.

More likely, the export of GALNTs-containing carriers results in the depletion of an unidentified receptor on Golgi membranes (Figure 7B). Indeed, unlike Arf-GTP, Arf-GDP requires a protein receptor to bind to Golgi membrane (Donaldson and Jackson, 2011). Two candidates have been proposed, the p23 protein and the SNARE membrin (Gommel *et al*., 2001; Honda *et al*., 2005). GBF1 also appears to require an additional receptor to bind efficiently to Golgi membranes (Quilty *et al*., 2018). It is possible that after Src activation, either the Arf-GDP or GBF1 receptors (or both) are transported from Golgi membranes by the tubules-derived carriers. This would explain why overnight Src expression, while inducing a marked GALNT relocation, also results in a reduction of Arf-GTP levels: the Arf receptor has been partially depleted at the Golgi. By contrast, wild-type GBF1 overexpression stably increased Arf-GTP levels. As GBF1 overexpression does not promote GALNTs relocation, there would not be with Arf-GDP on Golgi membranes. Mass spec data and residue conservation analysis suggest that Y876 and Y898 are particularly important for this effect, although the 8 other sites identified may also play a role. Y876 phosphorylation seems to induce the partial melting of an alpha-helix within the Sec7d, allowing for better binding to Arf. Partial melting is dependent on the phospho group interacting with either residues R843 or K844. Consistently, an R843A.K844A double mutant is unable to increase Arf1-GTP levels in response to Src activation. The R843E.K844E mutant is also unable to respond to Src, and in addition it induces a reduced basal Arf-GTP level. This is consistent with the Sec7d of this mutant being locked into a low Arf-binding conformation.

The effect of Y898 phosphorylation is less clear, it might release an hypothetical inhibitory interaction between the HDS1 domain and the Sec7d. The data suggest that phosphorylation on both residues synergise to stimulate binding of GBF1 to Arf1-GDP.

Y876 and Y898 residues are highly conserved among vertebrates and invertebrates homologues of GBF1, suggesting their phosphorylation is an ancient, conserved mechanism of regulation and could be involved in other processes. GBF1 is involved in different physiological situations in addition to the regulation of GALNTs activity (Kaczmarek, Verbavatz and Jackson, 2017). The axis Src-GBF1-Arf might also be involved in the regulation of Golgi organisation based on work by the Luini’s group(Pulvirenti *et al*., 2008; Consoli *et al*., 2012; Luini and Parashuraman, 2016). GBF1-Arf are also involved in the positioning of mitochondria, a locale where Src kinase has also been detected (Hebert-Chatelain, 2013; Ackema *et al*., 2014; Walch *et al*., 2018).

In addition, Y876 is conserved in all the Sec7d proteins we looked at. For several other GEFs, phosphorylation of the corresponding residue has been reported in various high-throughput studies. These findings suggest that the helix melting regulation is shared by other GEF proteins and could be a widespread mechanism for tyrosine kinases to regulate the GEF family of proteins and the associated membrane trafficking events.

To summarise, our study reveals a key mechanistic insight into how Src regulates Golgi to ER retrograde traffic. It also questions the classical model of membrane traffic between Golgi and in the GALA pathway might help interfere therapeutically with a process driving the invasiveness of solid tumor cancer cells.

## Materials and methods

### Cloning and cell culture

Wild type Hela cells were from V. Malhotra (CRG, Barcelona). HEK293T and A431 cells were a gift from W. Hong (IMCB, Singapore). NIH3T3 and NIH3T3-vsrc mouse fibroblast were a gift from X. Cao (IMCB, Singapore). All cells were maintained in DMEM with 10% fetal bovine serum (FBS) except for HEK293T which was cultivated in 15% FBS. All cells were grown at

37°C in a 10% CO_2_ incubator. Plasmids encoding full-length wild-type chicken SRC and an E378G mutant were a gift from Roland Baron (Harvard Medical School, Boston, MA). Src8A7F construct was obtained from Philip Cole’s laboratory (Johns Hopkins University School of Medicine). The human GalT-GFP construct corresponds to the first 81 AA of human beta

1,4-galactosyltransferase (GalT) fused in frame with Aequorea coerulescens green fluorescent protein, allowing targeting of the chimeric protein to medial and trans cisternae. The construct was purchased from Clontech Laboratories, Inc. Human GALNT2 (NM_004481) was cloned from a cDNA library generated from HT29 cells. All constructs were cloned into entry vector pDONR221 (Invitrogen, Life Technologies Corporation, Carlsbad, CA) and subsequently, gateway destination vectors expressing either emGFP or mcherry tag as described in (Gill *et al*.,

2010). All constructs were verified by sequencing and restriction enzyme digests before use. Hela and HEK293T cell lines stably expressing Src8A7F-Cmcherry or GALNT2-GFP were generated by lentiviral infection as described in (Gill *et al*., 2013) and subsequently, FACS sorted to enrich for mcherry- or GFP-expressing cells.

### Antibodies and reagents

*Helix pomatia* Lectin (HPL) conjugated with 647 nm fluorophore, Alexa Fluor secondary antibodies, and Hoechst 33342 were purchased from Invitrogen. Anti-GALNT1 for immunofluorescence staining was a gift from U. Mendel and H. Clausen (University of Copenhagen, Denmark). Anti-GBF1 antibody for immunoprecipitation was from BD Biosciences (Franklin Lakes, NJ). Anti-GBF1 (C-terminus), anti-Giantin and anti-Arf1 were from Abcam (Cambridge, MA). Human recombinant growth factors PDGF and EGF were purchased from BD Biosciences. Imidazole was purchased from Sigma-Aldrich (St. Louis, MO). GGA3 PBD agarose beads were purchased from Cell Biolabs, Inc. (San Diego, CA). GTP-trap agarose beads were purchased from ChromoTek GmbH, Germany.

### Automated image acquisition and quantification

The staining procedures were performed as described in (Chia *et al*., 2014). Briefly, images were acquired sequentially with a 20x objective on a laser scanning confocal high-throughput microscope (Opera Phenix™, PerkinElmer Inc.). Image analysis was performed using the Columbus Software (version 2.8.0). GFP and mcherry expressing cells were selected based on the intensity cutoff of the top 10% of expressing cells. The HPL staining intensity of the selected cell population was quantified by drawing a ring region outside the nucleus that covers most of the cell area. The HPL intensity per cell of each well quantified. Statistical significance was measured using a paired t test assuming a two-tailed Gaussian distribution.

### High-resolution fluorescence microscopy

The procedures were performed as described in (Chia *et al*., 2014). Briefly, cells were seeded onto glass coverslips in 24-well dishes (Nunc, Denmark) before various treatments. They were fixed with 4% paraformaldehyde-4% sucrose in D-PBS, permeabilized with 0.2% Triton-X for 10 minutes and stained with the appropriate markers. This was followed by secondary antibody staining for 20 minutes before mounting onto glass slides using FluorSave (Merck)). The cells imaged at room temperature using an inverted confocal microscope (IX81; Olympus Optical Co. Ltd, Tokyo, Japan) coupled with a CCD camera (model FVII) either with a 60× objective (U Plan Super Apochromatic [UPLSAPO]; NA 1.35) or 100× objective (UPLSAPO; NA 1.40) under Immersol oil. Images were processed using Olympus FV10-ASW software.

### High-resolution live imaging

For imaging of GALNT tubules upon imidazole treatment, cells were seeded on 8-chamber glass chambers (Thermoscientific, #155411) and treated with 5 mM imidazole.

For the PDGF stimulation, cells were seeded in 6-channel μ-Slide slides (ibidi GmbH, Germany) and treated with 50ng/ml of PDGF stimulation using a perfusion pump system (ibidi GmbH) to inject the media at a constant and gentle flow rate. The cells placed in a 37°C environmental chamber and imaged using an inverted confocal microscope (IX81; Olympus Optical Co. Ltd, Tokyo, Japan) coupled with a CCD camera (model FVII) with a 100× objective (UPLSAPO; NA 1.40) under Immersol oil. Images were processed using Olympus FV10-ASW software.

### Immunoprecipitation (IP) and western blot analysis

Procedures for cell harvesting and processing for IP and western blot were performed as described previously with some modifications (Gill *et al*., 2010). For imidazole treatment and growth factor stimulations, cells were serum starved for 24 hours before treatment with 5 mM imidazole, 50ng/ml PDGF or 100ng/ml EGF respectively. Cells were washed twice using ice-cold D-PBS before scraping in ice-cold RIPA lysis buffer (50 mM Tris [pH 7.4], 150 mM NaCl, 1% NP-40 alternative, complete protease inhibitor and phosphatase inhibitor [Roche Applied Science, Mannheim, Germany]). The lysate was incubated on ice for 30 minutes with gradual agitation before clarification of samples by centrifugation at 10,000 ×g for 10 minutes at 4°C. Clarified lysate protein concentrations were determined using Bradford reagent (Bio-Rad Laboratories, Hercules, CA) before sample normalisation. To IP endogenous GBF1, samples were incubated with 2.5μg of GBF1 (BD Biosciences) for one hour at 4°C with constant mixing. The IP samples were then incubated with 20μl of washed protein A/G–Sepharose beads (Millipore) for two hours at 4°C with constant mixing. IP samples were washed three times with 1 ml of RIPA buffer containing complete protease inhibitor and phosphatase inhibitor. For NGFP-GBF1 IP, clarified cell lysates were incubated with GTP-trap agarose beads (ChromoTek GmbH, Germany) for two hours before washing with GFP wash buffer (10 mM Tris [pH 7.5], 150 mM NaCl, 0.5 mM EDTA, complete protease inhibitor and phosphatase inhibitor) for three times. For Arf1-GTP loading assay, the clarified cell lysates were incubated with GGA3 PBD agarose beads (Cell Biolabs Inc, CA) for one hour at 4°C with agitation before washing. Samples were diluted in lysis buffer with 4× SDS loading buffer and boiled at 95°C for ten minutes. The proteins were resolved by SDS-PAGE electrophoresis using bis-tris NuPage gels (Invitrogen) and transferred to PVDF or nitrocellulose membranes. Membranes were then blocked using 3% BSA dissolved in Tris-buffered saline with tween (TBST: 50 mM Tris [pH 8.0, 4°C], 150 mM NaCl, and 0.1% Tween 20) for one hour at room temperature before incubation with primary antibodies overnight. Membranes were washed three times with TBST before incubation with secondary HRP-conjugated antibodies (GE Healthcare). Membranes were further washed three times with TBST before ECL exposure.

### In vitro Arf1 binding assay

Procedures for cell harvesting and processing for IP and western blot were described as above. NGFP-GBF1 expressed in cells was IP with GTP-trap agarose beads and washed three times with GFP wash buffer in the presence of complete protease inhibitor and once with HKMT buffer (20 mM HEPES, pH 7.4, 0.1 M KCl, 1 mM MgCl2, 0.5% Triton X-100) containing complete protease inhibitor and phosphatase inhibitor. The purified NGFP-GBF1 on agarose beads were then incubated with either 4mg of cell lysates of HEK293T cells expressing Arf1-V5 (agarose beads pre-cleared) or 10ug recombinant Arf1-del17 protein for one hour for 4°C. Subsequently, the beads were washed three times with the HKMT buffer to remove unbound Arf1 protein. The beads were boiled at 95°C for ten minutes. The amount of GBF1 bound Arf1 was resolved by western blotting.

### LC/MS analysis

The GFP-GBF1 bands of immunoprecipitated samples run on a SDS-PAGE using a NuPAGE 4-12% Bis Tris Gel (Invitrogen) were excised followed by in-gel digestion as described previously (Shevchenko *et al*., 2006). The peptide samples were subjected to a LTQ Orbitrap classic for data dependent acquisition and a Q-Exactive for parallel reaction monitoring (Thermo Fisher Scientific) analysis as described previously (Swa *et al*., 2012).

For parallel reaction monitoring (PRM) on Q-Exactive, targeted MS2 was carried out using a resolution of 17,500, target AGC values of 2E5 with maximum injection time of 250 ms, isolation windows of 2 Th and a normalized collision energy of 27. MS/MS scans started from m/z 100.

### Data processing and database search

Raw file obtained from data dependent acquisition was processed using Mascot Daemon (version 2.3.2, Matrix Science). Data import filter for precursor masses from 700 to 4000 Da, with a minimum scans per group of 1 and a minimum peak count of 10. Mascot search was performed using the IPI Human database (ipi.HUMAN.v3.68.decoy.fasta or ipi.HUMAN.v3.68.decoy.fasta), trypsin as enzyme and two allowed missed cleavages. Carbamidomethyl (C) was set as a static modification while the dynamic modifications were Acetyl (Protein N-term), Oxidation (M) and Phosphorylation (S/T/Y). Tolerance for the precursor masses was 7ppm and for fragments 0.5 Da for samples analysed on LTQ Orbitrap. Raw file obtained from parallel reaction monitoring was processed using open-source Skyline software tool [Maclean, B. *et al*. Bioinformatics 2010, 26, 966], (http://skyline.maccosslab.org.). The accuracy of the peaks assigned by Skyline was manually validated using Thermo Xcalibur Qual Browser by manual inspection of the targeted MS2 spectra and by XIC to ensure the m/z of the fragment ions are within 20 ppm of their theoretical values.

### Structure modelling and molecular dynamics

The 3D structure of GBF1 protein is not available, therefore a structural model of the Sec7 domain of GBF1 (GBF1_Sec7) protein was generated using comparative modeling methods (Sali and Blundell, 1993). Homology model of the GBF1_Sec7 in its autoinhibited form was generated using the crystal structure of the autoinhibited form of Grp1 Arf GTPase Exchange Factor (PDB: 2R0D, resolution 2.0 Å) which shares ∼65% homology with GBF1 in the Sec7 domain. A 3D structural model of the GBF1_Sec7-Arf1 complex was generated using the crystal structure of Arno_Sec7-Arf1 (PDB: 1R8Q, resolution 1.9 Å) since Arno shares ∼65% homology with GBF1 in the Sec7 domain.

MD simulations were carried out with the pemed.CUDA module of the program Amber18 (Case *et al*., no date) using standard and well tested protocols (Kannan *et al*., 2015). All atom versions of the Amber 14SB force field (ff14SB) (Maier *et al*., 2015) were used to represent the protein. Force field parameters for phosphorylated tyrosine and GTP were taken as described elsewhere (Homeyer *et al*., 2006); An overall charge of -2e is assigned to the phosphate groups. The Xleap module was used to prepare the system for the MD simulations. All the simulation systems were neutralized with appropriate numbers of counterions. Each neutralized system was solvated in an octahedral box with TIP3P (Jorgensen *et al*., 1983) water molecules, leaving at least 10 Å between the solute atoms and the borders of the box. All MD simulations were carried out in explicit solvent at 300K. During the simulations, the long-range electrostatic interactions were treated with the particle mesh Ewald (Darden, York and Pedersen, 1993) method using a real space cutoff distance of 9 Å. The Settle (Miyamoto and Kollman, 1992) algorithm was used to constrain bond vibrations involving hydrogen atoms, which allowed a time step of 2 fs during the simulations. Solvent molecules and counterions were initially relaxed using energy minimization with restraints on the protein and inhibitor atoms. This was followed by unrestrained energy minimization to remove any steric clashes. Subsequently the system was gradually heated from 0 to 300 K using MD simulations with positional restraints (force constant: 50 kcal mol-1 Å-2) on the protein atoms over a period of 0.25 ns allowing water molecules and ions to move freely. During an additional 0.25 ns, the positional restraints were gradually reduced followed by a 2 ns unrestrained MD simulation to equilibrate all the atoms. Production runs were carried out for 250 ns in triplicates (assigning different distributions of initial velocities) for each system. Simulation trajectories were visualized using VMD (Humphrey, Dalke and Schulten, 1996) and figures were generated using Pymol.

Binding free energies and per-residue decomposition of binding free energies between the GBF1_Sec7 (unphosphorylated and phosphorylated at Tyr876) and Arf1 were calculated using the standard MMPBSA approach (Hou *et al*., 2011; Homeyer and Gohlke, 2012). Conformations extracted from the last 125 ns of the MD simulations of each GBF1_Sec7-Arf1 complex were used and binding energy calculations/per residue decomposition analysis were carried out using standard protocols (Kannan *et al*., 2015). Buried Surface Area (BSA) was computed using the program NACCESS(Hubbard and Thornton, 1993) .

## Supplementary Figures

Supplementary Figure 1

(A) Schematic of imidazole (imdz) rescue of Src8A7F mutant in comparison to wild type Src. (B) SDS-PAGE comparison of the total phosphotyrosine levels of imdz treated Hela-IS cells over time and cells expressing empty mcherry vector, SrcKM and SrcEG mutants. (C) Images of Src8A7F expression as well as HPL and Golgi marker Giantin staining of the cells shown in Figure 1B over time of imdz stimulation. Images were acquired under constant acquisition settings using an automated confocal microscope. Scale bar: 20 μm. (D) HPL staining of Hela cells expressing inactive SrcKM and active SrcEG mutants. (E) Quantification of HPL levels over duration of 5 mM imdz stimulation in wildtype Hela cells. Values were normalised with respect to untreated cells (0h). (F) HPL staining of Hela-IS stable cell line over time of imdz washout. Cells were treated with 5 mM imdz for 24 hours prior to washout. Scale bar: 50 μm. (G) Quantification of HPL levels over duration of imdz treatment (blue bars) and washout of imdz and fixed over various durations after 24 hours of imdz treatment (green bars). Values were normalized with respect to untreated cells (0h). (H) HPL staining of Hela cells after 50ng/ml PDGF stimulation. Scale bar: 10 μm. (I) Quantification of HPL levels after PDGF stimulation normalized with respect to untreated cells (0h). (J) Stills of the movie demonstrating GALNT2 tubule formation in Hela cells stimulated with 50ng/ml PDGF. Scale bar: 5 μm. (K) The Golgi glycosyltransferase GalT was not observed in the GALNT2 tubules. Scale bar: 10 μm. (L) Images of BCOP localisation of Hela-IS cells over time with imdz treatment. (M) Images of BCOP localisation acquired from automated microscope and used for quantification. GM130 is a Golgi marker. Scale bar: 20 μm. (N) Quantification of levels BCOP at the Golgi over time of imdz treatment. Values on graphs indicate the mean ± SD. Statistical significance (p) measured by two-tailed paired t test.*, p < 0.05, **, p<0.01 and ***, p < 0.001 relative to untreated cells. NS, non significant.

Supplementary Figure 2

(A) Additional representative images of Arf1 on GALNT2 tubules emanating from the Golgi upon 10 minutes stimulation of 20mM imdz. Images were acquired at 100x magnification under immersol oil. Scale bar: 5 μm. (B) SDS-PAGE analysis of the levels of Arf1-GTP IP using GGA3 beads in HEK293T cells expressing empty mcherry vector, SrcKM and SrcEG mutants. (C) Quantification of the levels of Arf1-GTP in (B). Three experimental replicates were measured. (D) SDS-PAGE analysis of total lysate (L), cytoplasmic (C) and membrane (M) levels of various proteins after subcellular fractionation. ER resident Calnexin (CANX), Golgi marker GM130 as well as cytoplasmic actin and MAP kinase ERK8 were shown. (E) siRNA knockdown efficiencies of various proteins assayed. siNT refers to non-targeting siRNA. Values on graphs indicate the mean ± SD. Statistical significance (p) measured by two-tailed paired t test.*, p < 0.05 and **, p<0.001 relative to untreated cells. NS, non significant.

Supplementary Figure 3

(A) HPL staining of Hela-IS stable cell line treated with siRNA targeting GBF1 before and after 4 hours of imdz treatment. siNT refers to non-targeting siRNA and siGALNT1+T2 refers to co-transfection of GALNT1 and GALNT2 siRNAs. Images were acquired under constant acquisition settings using an automated confocal microscope. Scale bar: 50 μm. (B) Quantification of HPL staining intensity per cell normalized to the respective untreated cells (0h) for each siRNA treatment. (C) Immunoblot analysis of the levels of Tn modified ER resident PDIA3 from VVL IP in HEK-IS cell line upon GBF1 siRNA knockdown. Cells were untreated or treated with 5 mM imdz for 6 hours. (D) Schematic illustrating the workflow of the in vitro Arf1 binding assay. (E) Quantification of the levels of bound Arf1-V5 to GFP and GFP-GBF1 (WT) IP from cells expressing inactive SrcKM or active SrcEG in the in vitro binding assay shown in Figure 3G. Results representative of two experimental replicates. (F) SDS-PAGE analysis of the levels of recombinant protein Arf1-del17-His bound to GFP-GBF1 IP from inactive SrcKM or active SrcEG expressing cells in an in vitro binding assay. (G) Images from time lapse imaging of GALNT2-GFP in Hela-IS cells that were either treated with siRNA targeting GBF1 (siGBF1) or siNT stimulated with 5 mM imdz. Scale bar: 5 μm. (H) Quantification of the number of tubules observed in the first 30mins upon imidazole treatment. Values on graphs indicate the mean ± SD. Statistical significance (p) measured by two-tailed paired t test.*, p < 0.05 and **, p<0.01 relative to untreated (0h) or GFP expressing cells. NS, non significant.

Supplementary Figure 4

(A) SDS-PAGE analysis of Y876 phosphorylation levels in endogenous GBF1 in cells expressing empty mcherry vector, inactive SrcKM or active SrcEG. (B) Immunoblot analysis of the levels of Tn modified PDIA4 from VVL IP in mouse embryonic fibroblasts WT, SYF and SYFsrc as well as mouse fibroblasts NIH3T3 WT and 3T3vSrc. SYF cells are knockout of Src, Yes and Fyn while SYFsrc cells are SYFcells with stable transfection of c-Src. 3T3vsrc cells are v-Src transformed 3T3 cells. (C) Corresponding coomassie staining of immunoprecipitated GFP and GFP-GBF1 purified from HEK293T cells that were used for in vitro Src kinase assay. The purified proteins on the beads were incubated with recombinant Src protein in the presence or absence of nucleotide ATP. (D) Table of the mascot scores and the frequencies of peptide-spectrum matches (PSM) that are more than or equal to 15 for each phosphosite on GBF1 that is co-expressed with SrcKM (KM) or SrcEG (EG). GBF1 was cleaved with either trypsin or endoproteinase AspN for analysis. (E) Table illustrating the conservation of each identified tyrosine residues that were found to be phosphorylated by Src. (F) Quantification of the peak area of the SILAC mass spectral of the peptides containing Y876 (DFEQDILEDMyHAIK) and Y898 (ENyVWNVLLHR) phosphorylation in SrcKM (blue bars) or SrcEG (red bars). (G) Mass spectra of Y876 phosphopeptide. (H) Mass spectra of Y898 phosphopeptide. (I) SDS-PAGE analysis of total pY levels on endogenous GBF1 in Hela cells expressing empty mcherry vector, inactive SrcKM or active SrcEG. GBF1 was IP with an antibody targeting the N-terminus of the protein. (J) SDS-PAGE analysis of Y876 phosphorylation on GFP-GBF1 IP from HEK-IS cell line over various durations of imidazole treatment. (K) In vitro phosphorylation assay of GFP and GFP-GBF1 wild type or mutant with recombinant Src protein. Total phosphorylation and phosphorylation of GBF1 at Y876 is detected by pY(4G10) and 2P4 antibodies respectively.

Supplementary Figure 5

(A) SDS-PAGE analysis of total Arf1 and GTP loaded Arf1 in HEK293T cells expressing various mutants of GBF1. Y876E and Y898E are phospho-mimetic mutants while Y876F and Y898F are phospho-null mutants. Two experimental replicates for each condition were shown in the blot. (B) Quantification of Arf1-GTP loading in (A). (C) Representative images of HPL staining in Hela cells co-expressing wild type GBF1 or phospho-null mutants with active SrcEG or inactive SrcKM. Scale bar: 50 μm. (D) Quantification of HPL staining levels of Hela-IS cells co-expressing with wild type or mutant GBF1 without (blue bars) or with 4 hours imdz treatment (pink bars). Values were from three experimental replicates. (E) SDS-PAGE analysis of the levels of recombinant Arf1-His bound to wild type or mutant GFP-GBF1 IP from cells expressing inactive SrcKM or active SrcEG in an *in vitro* binding assay. Values on graphs indicate the mean ± SD. Statistical significance (p) measured by two-tailed paired t test.*, p < 0.05 and **, p<0.001 relative to untreated (0h) or GFP expressing cells. NS, non significant.

Supplementary Figure 6

(A) Electrostatic map of the charged residues on GBF1 Sec7d in the presence (right) and absence (left) of phosphorylation on Y898 on the C-terminal linker. (B) MD snapshot of the release of the C-terminal linker from the main body of the Sec7d when Y898 is phosphorylated. Values on graphs indicate the mean ± SD. Statistical significance (p) measured by two-tailed paired t test.*, p < 0.05 and **, p < 0.001 relative to untreated cells or to 10-min imdz treated cells expressing wild type GBF1. NS, non significant.

Supplementary Figure 7

(A) Representation of the predicted electrostatic bonds between the GBF1: Arf1 complex in the unphosphorylated (WT) and Y876 phosphorylated states. The Sec7d was shown in blue while Arf1 protein in grey. Residues involved in inter-protein interactions are shown as sticks, with inter-protein H-bonds highlighted as black dashed lines. (B) Per-residue decompositions of the binding free energies of interactions between the Y876-phosphorylated GBF1 Sec7d and Arf1 and between unphosphorylated GBF1 Sec7d and Arf1, using the MMGBSA approximations averaged over the conformations generated from MD simulations of the complexes. (C) Representative images of HPL staining in Hela cells co-expressing wild type or HI loop mutants with active SrcEG. Scale bar: 50 μm. (D) Quantification of HPL staining levels of cells co-expressing wild type or mutant GBF1 with inactive SrcKM (blue bar) or active SrcEG (pink bars). Values were from six replicates from two independent experiments. (E) Representative images of HPL staining in Hela-IS cells co-expressing wild type or HI loop mutants. Scale bar: 50 μm. (F) Quantification of HPL staining levels of Hela-IS cells co-expressing wild type or mutant GBF1 that were unstimulated (0h, blue bar) or stimulated with 5 mM imdz (4h, pink bars). Values were from three replicates. Values on graphs indicate the mean ± SEM. Statistical significance (p) measured by two-tailed paired t test.*, p < 0.05 and **, p<0.001 relative to control cells. NS, non significant.

