## Supplementary material for "Src activates retrograde membrane traffic through phosphorylation of GBF1": Supp figure S1-7

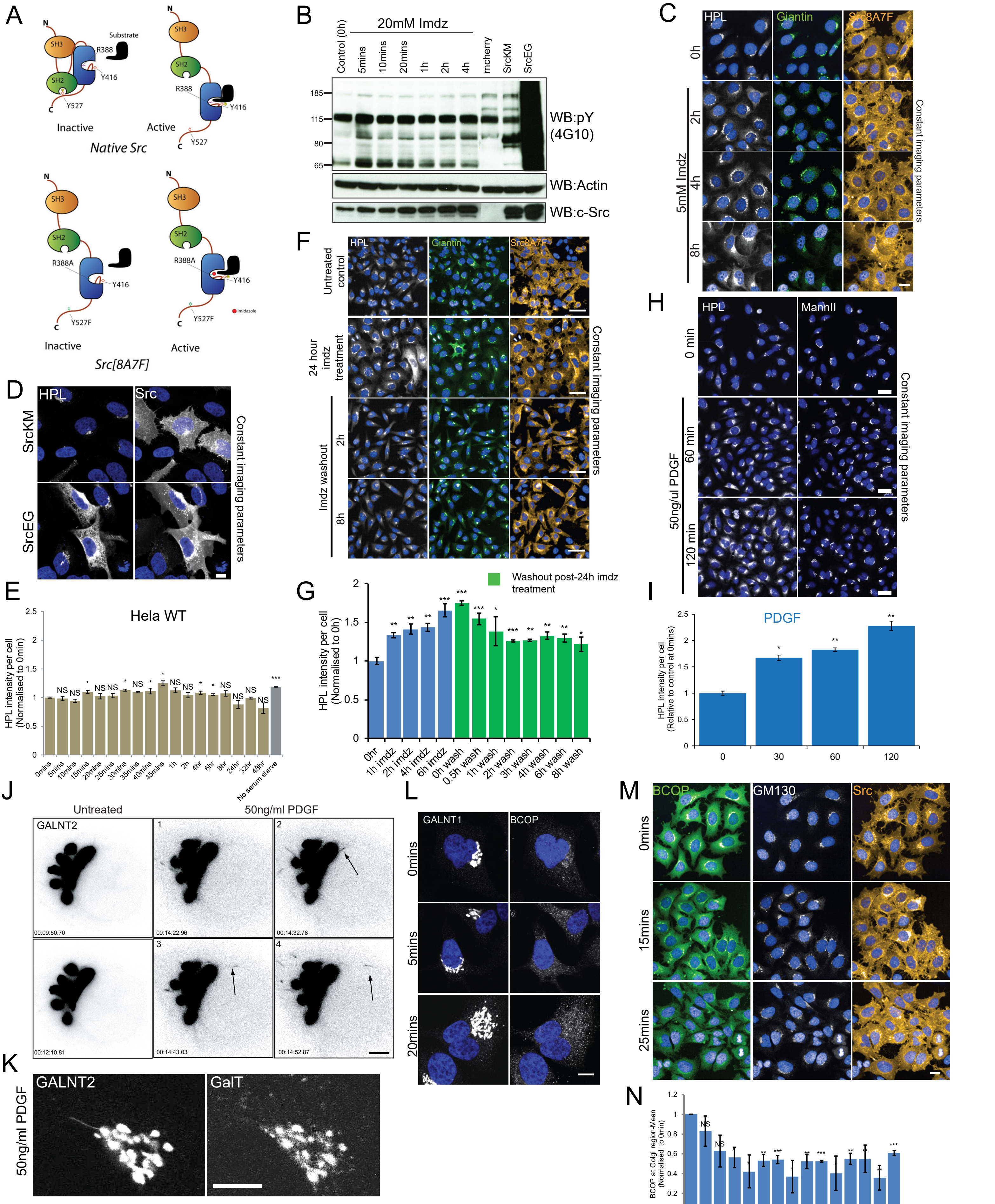

Supplementary Figure 1

(A) Schematic of imidazole (imdz) rescue of Src8A7F mutant in comparison to wild type Src. (B) SDS-PAGE comparison of the total phosphotyrosine levels of imdz treated Hela-IS cells over time and cells expressing empty mcherry vector, SrcKM and SrcEG mutants. (C) Images of Src8A7F expression as well as HPL and Golgi marker Giantin staining of the cells shown in Figure 1B over time of imdz stimulation. Images were acquired under constant acquisition settings using an automated confocal microscope. Scale bar: 20 µm. (D) HPL staining of Hela cells expressing inactive SrcKM and active SrcEG mutants. (E) Quantification of HPL levels over duration of 5 mM imdz stimulation in wildtype Hela cells. Values were normalised with respect to untreated cells (0h). (F) HPL staining of Hela-IS stable cell line over time of imdz washout. Cells were treated with 5 mM imdz for 24 hours prior to washout. Scale bar: 50 µm. (G) Quantification of HPL levels over duration of imdz treatment (blue bars) and washout of imdz and fixed over various durations after 24 hours of imdz treatment (green bars). Values were normalized with respect to untreated cells (0h). (H) HPL staining of Hela cells after 50ng/ml PDGF stimulation. Scale bar: 10 µm. (I) Quantification of HPL levels after PDGF stimulation normalized with respect to untreated cells (0h). (J) Stills of the movie demonstrating GALNT2 tubule formation in Hela cells stimulated with 50ng/ml PDGF. Scale bar: 5 µm. (K) The Golgi glycosyltransferase GalT was not observed in the GALNT2 tubules. Scale bar: 10 µm. (L) Images of BCOP localisation of Hela-IS cells over time with imdz treatment. (M) Images of BCOP localisation acquired from automated microscope and used for quantification. GM130 is a Golgi marker. Scale bar: 20 µm. (N) Quantification of levels BCOP at the Golgi over time of imdz treatment. Values on graphs indicate the mean ± SD. Statistical significance (p) measured by two-tailed paired t test.*, p < 0.05, **, p<0.01 and ***, p < 0.001 relative to untreated cells. NS, non significant.

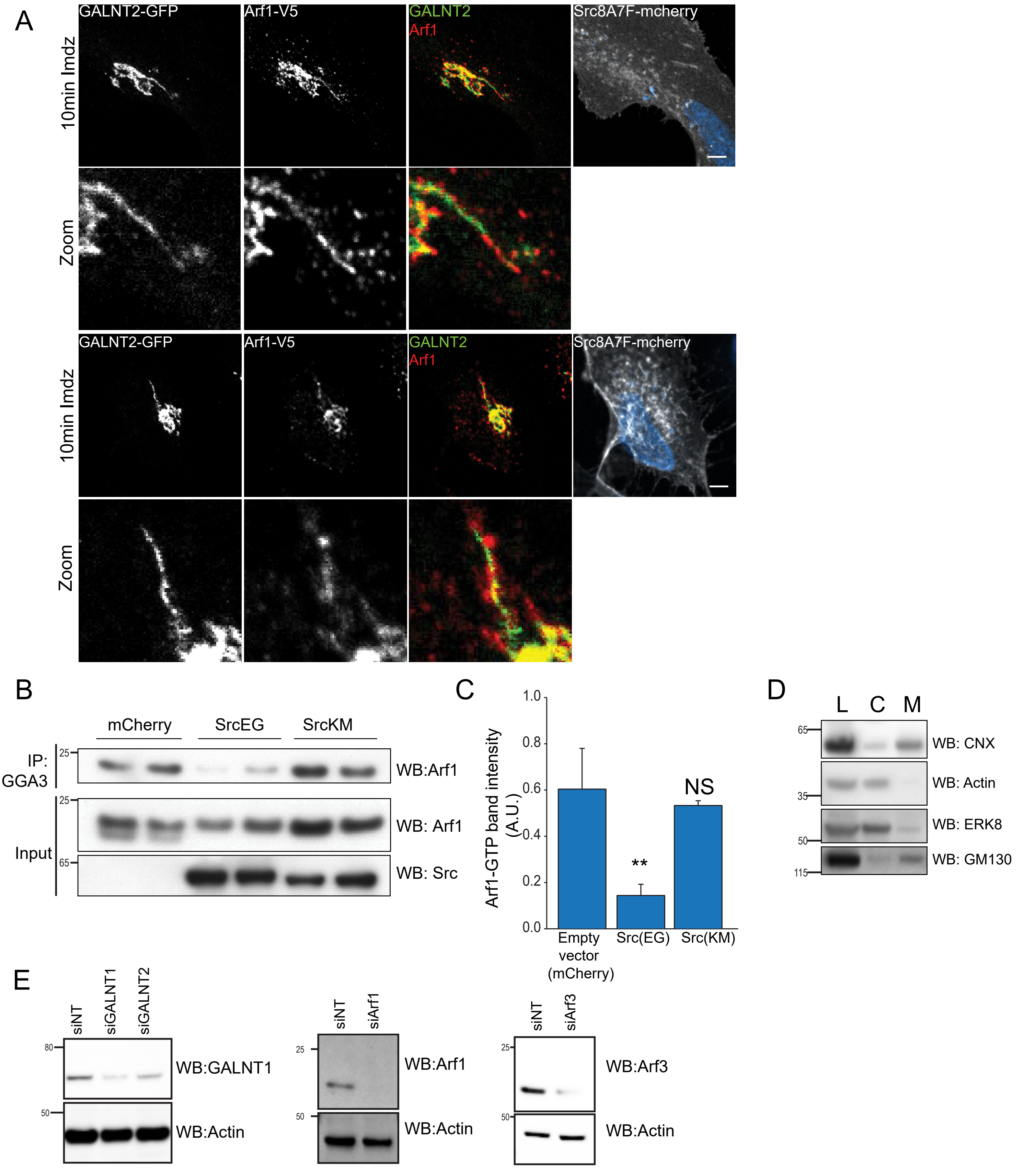


Supplementary Figure 2

(A) Additional representative images of Arf1 on GALNT2 tubules emanating from the Golgi upon 10 minutes stimulation of 20mM imdz. Images were acquired at 100x magnification under immersol oil. Scale bar: 5 µm. (B) SDS-PAGE analysis of the levels of Arf1-GTP IP using GGA3 beads in HEK293T cells expressing empty mcherry vector, SrcKM and SrcEG mutants. (C) Quantification of the levels of Arf1-GTP in (B). Three experimental replicates were measured. (D) SDS-PAGE analysis of total lysate (L), cytoplasmic (C) and membrane (M) levels of various proteins after subcellular fractionation. ER resident Calnexin (CNX), Golgi marker GM130 as well as cytoplasmic actin and MAP kinase ERK8 were shown. (E) siRNA knockdown efficiencies of various proteins assayed. siNT refers to non-targeting siRNA. Values on graphs indicate the mean ± SD. Statistical significance (p) measured by two-tailed paired t test.*, $p < 0.05$ and **, $p<0.001$ relative to untreated cells. NS, non significant.

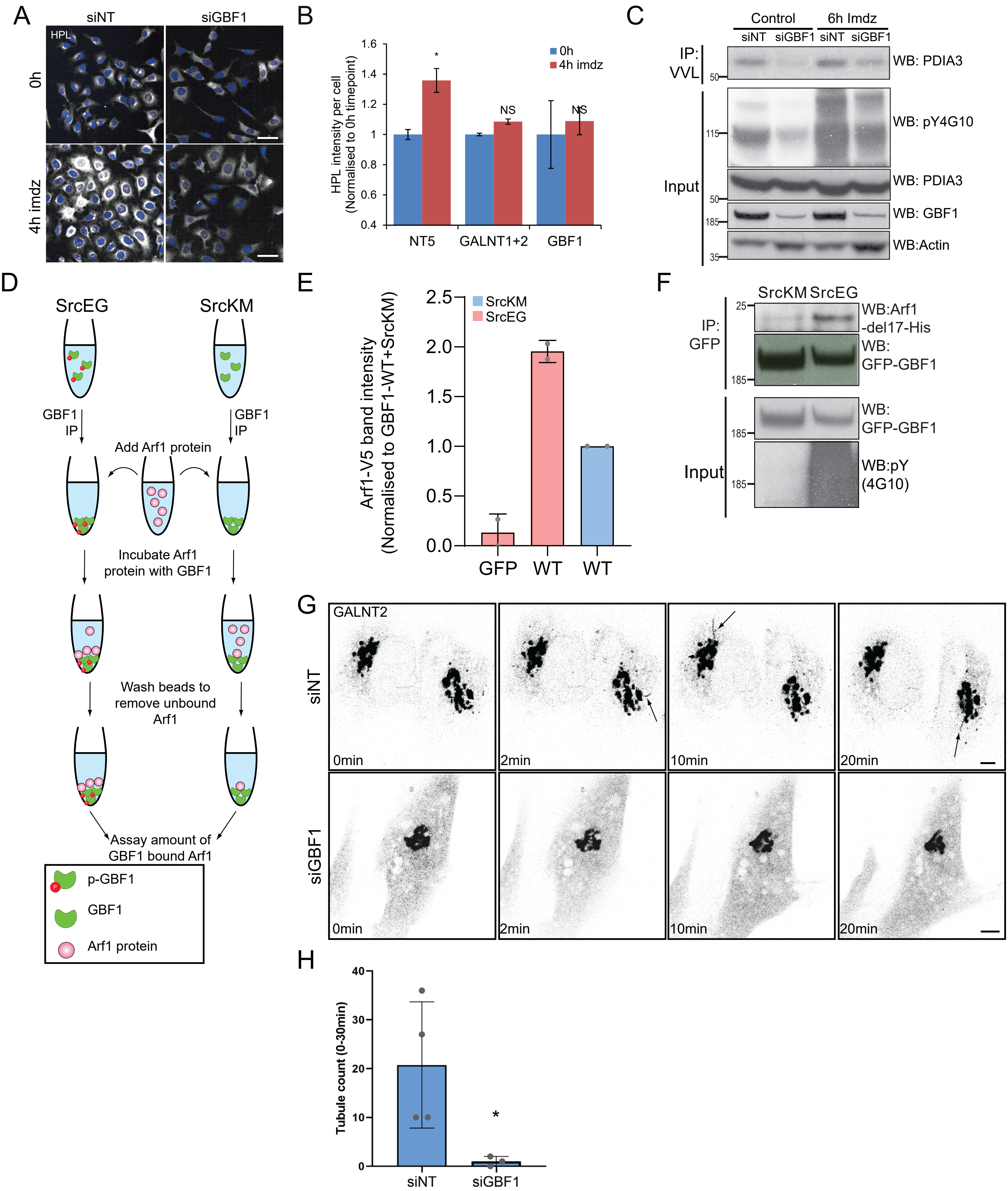


Supplementary Figure 3

(A) HPL staining of Hela-IS stable cell line treated with siRNA targeting GBF1 before and after 4 hours of imdz treatment. siNT refers to non-targeting siRNA and siGALNT1+T2 refers to co-transfection of GALNT1 and GALNT2 siRNAs. Images were acquired under constant acquisition settings using an automated confocal microscope. Scale bar: 50 μm. (B) Quantification of HPL staining intensity per cell normalized to the respective untreated cells (0h) for each siRNA treatment. (C) Immunoblot analysis of the levels of Tn modified ER resident PDIA3 from VVL IP in HEK-IS cell line upon GBF1 siRNA knockdown. Cells were untreated or treated with 5 mM imdz for 6 hours. (D) Schematic illustrating the workflow of the in vitro Arf1 binding assay. (E) Quantification of the levels of bound Arf1-V5 to GFP and GFP-GBF1 (WT) IP from cells expressing inactive SrcKM or active SrcEG in the in vitro binding assay shown in Figure 3G. Results representative of two experimental replicates. (F) SDS-PAGE analysis of the levels of recombinant protein Arf1-del17-His bound to GFP-GBF1 IP from inactive SrcKM or active SrcEG expressing cells in an in vitro binding assay. (G) Images from time lapse imaging of GALNT2-GFP in Hela-IS cells that were either treated with siRNA targeting GBF1 (siGBF1) or siNT stimulated with 5 mM imdz. Scale bar: 5 μm. (H) Quantification of the number of tubules observed in the first 30mins upon imidazole treatment. Values on graphs indicate the mean ± SD. Statistical significance (p) measured by two-tailed paired t test.*, $p < 0.05$ and **, $p<0.01$ relative to untreated (0h) or GFP expressing cells. NS, non significant.

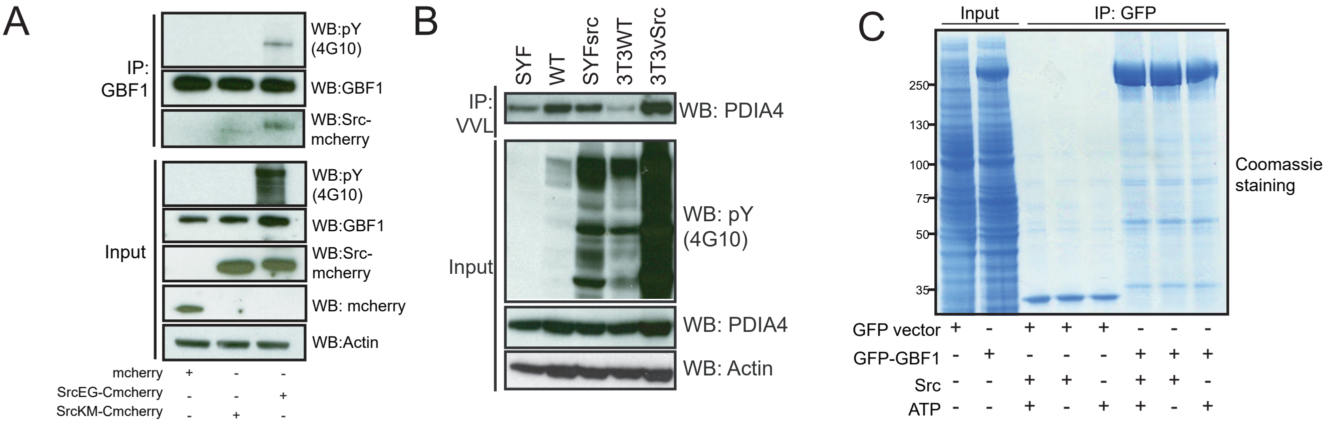


D

| Tyr position | Mascot Score | | Enzyme | #PSM for each phosphosite score ≥ 15 in kinase activated (Trypsin digestion) | #PSM for each phosphosite score ≥ 15 in kinase activated (AspN digestion) |
|---|---|---|---|---|---|
| | EG | KM | | | |
| 317 | 48 | N.D. | AspN | - | 3 |
| 377 | 51 | N.D. | AspN | - | - |
| 515 | 17 | 14 | Trypsin | 1 | - |
| 876 | 41 | N.D. | Trypsin | 20 | 7 |
| 898 | 33 | N.D. | Trypsin | 1 | - |
| 1244 | 41 | N.D. | Trypsin | 3 | 3 |
| 1373 | 28 | N.D. | AspN | 12 | 4 |
| 1487 | 46 | N.D. | AspN | - | 4 |
| 1513 | 91 | N.D. | Trypsin | 6 | - |
| 1567 | 20 | 1 | Trypsin | 1 | - |

E

| Tyr position | H. sapiens | M. Musculus | S. cerevisae | C.elegans | D.rerio |
|---|---|---|---|---|---|
| 317 | Y | - | - | - | - |
| 377 | Y | Y | - | - | Y |
| 515 | Y | Y | - | - | Y |
| 876 | Y | Y | Y | Y | Y |
| 898 | Y | Y | - | Y | Y |
| 1244 | Y | Y | - | - | Y |
| 1373 | Y | Y | - | - | Y |
| 1487 | Y | Y | - | - | Y |
| 1513 | Y | Y | - | - | Y |
| 1567 | Y | Y | - | - | Y |

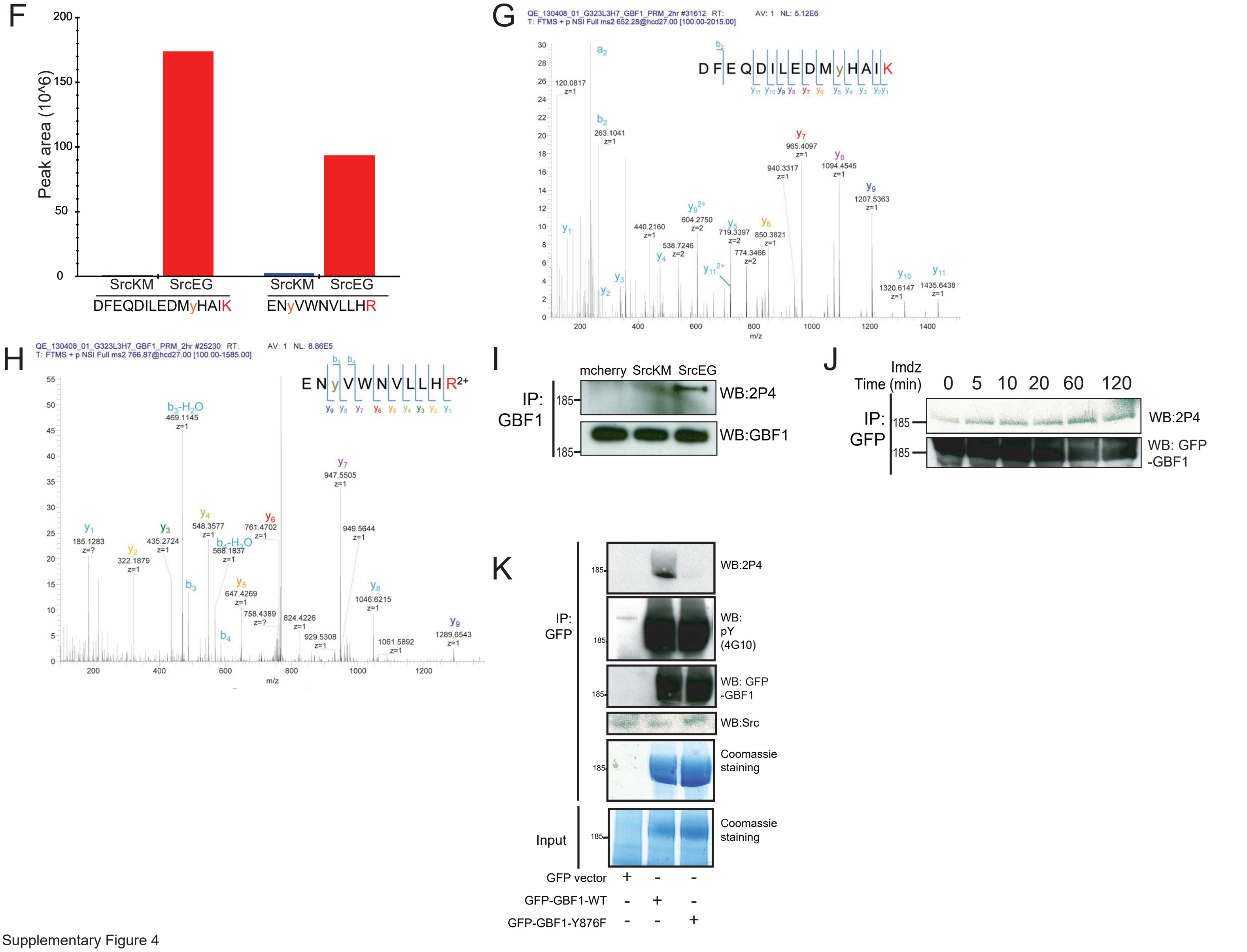


Supplementary Figure 4

(A) SDS-PAGE analysis of Y876 phosphorylation levels in endogenous GBF1 in cells expressing empty mcherry vector, inactive SrcKM or active SrcEG. (B) Immunoblot analysis of the levels of Tn modified PDIA4 from VVL IP in mouse embryonic fibroblasts WT, SYF and SYFsrc as well as mouse fibroblasts NIH3T3 WT and 3T3vSrc. SYF cells are knockout of Src, Yes and Fyn while SYFsrc cells are SYFcells with stable transfection of c-Src. 3T3vsrc cells are v-Src transformed 3T3 cells. (C) Corresponding coomassie staining of immunoprecipitated GFP and GFP-GBF1 purified from HEK293T cells that were used for in vitro Src kinase assay. The purified proteins on the beads were incubated with recombinant Src protein in the presence or absence of nucleotide ATP. (D) Table of the mascot scores and the frequencies of peptide-spectrum matches (PSM) that are more than or equal to 15 for each phosphosite on GBF1 that is co-expressed with SrcKM (KM) or SrcEG (EG). GBF1 was cleaved with either trypsin or endoproteinase AspN for analysis. (E) Table illustrating the conservation of each identified tyrosine residues that were found to be phosphorylated by Src. (F) Quantification of the peak area of the SILAC mass spectral of the peptides containing Y876 (DFEQDILEDMyHAIK) and Y898 (ENyVWNVLLHR) phosphorylation in SrcKM (blue bars) or SrcEG (red bars). (G) Mass spectra of Y876 phosphopeptide. (H) Mass spectra of Y898 phosphopeptide. (I) SDS-PAGE analysis of total pY levels on endogenous GBF1 in Hela cells expressing empty mcherry vector, inactive SrcKM or active SrcEG. GBF1 was IP with an antibody targeting the N-terminus of the protein. (J) SDS-PAGE analysis of Y876 phosphorylation on GFP-GBF1 IP from HEK-IS cell line over various durations of imidazole treatment. (K) In vitro phosphorylation assay of GFP and GFP-GBF1 wild type or mutant with recombinant Src protein. Total phosphorylation and phosphorylation of GBF1 at Y876 is detected by pY(4G10) and 2P4 antibodies respectively.

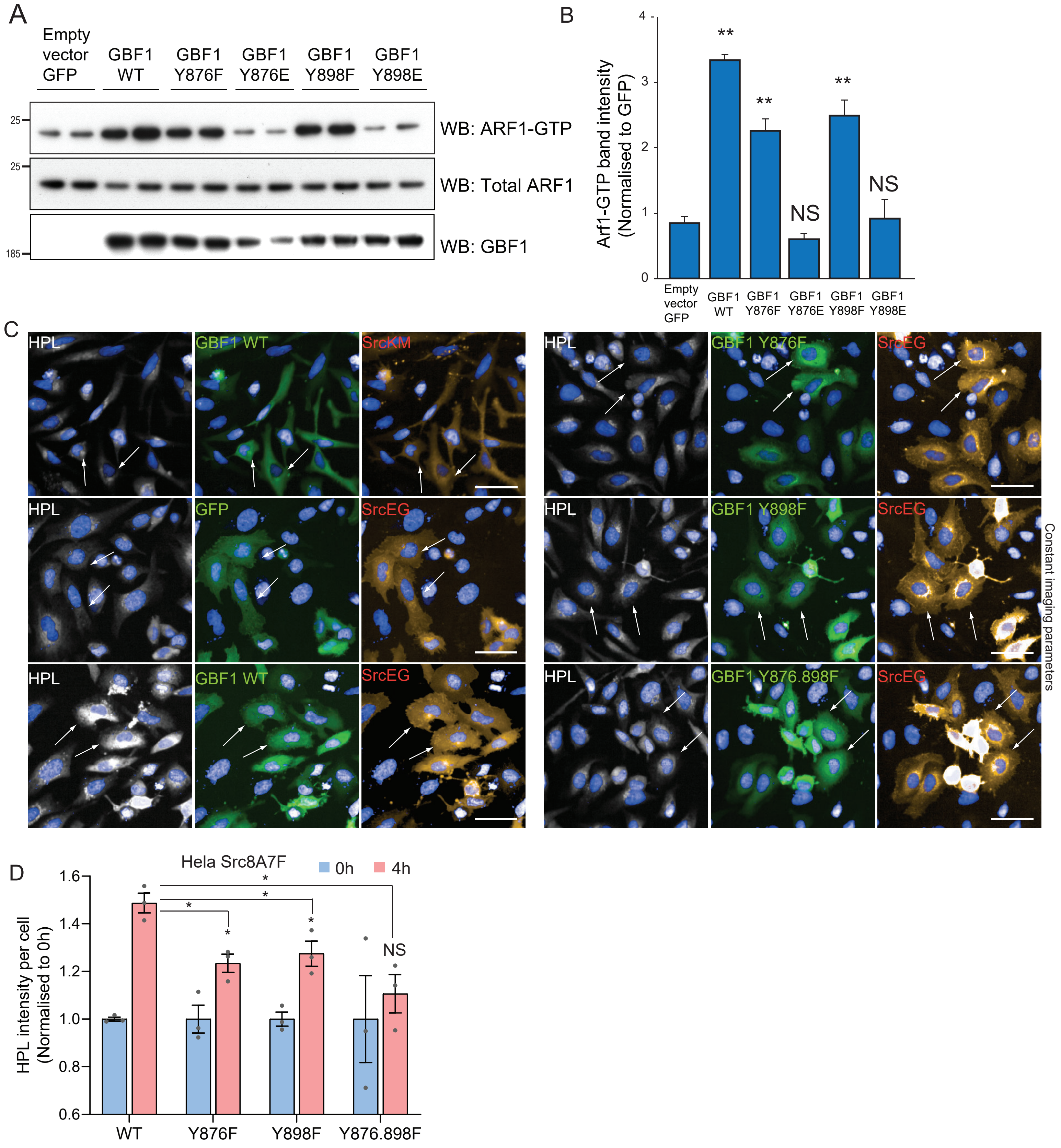


Supplementary Figure 5

(A) SDS-PAGE analysis of total Arf1 and GTP loaded Arf1 in HEK293T cells expressing various mutants of GBF1. Y876E and Y898E are phospho-mimetic mutants while Y876F and Y898F are phospho-null mutants. Two experimental replicates for each condition were shown in the blot. (B) Quantification of Arf1-GTP loading in (A). (C) Representative images of HPL staining in Hela cells co-expressing wild type GBF1 or phospho-null mutants with active SrcEG or inactive SrcKM. Scale bar: 50 µm. (D) Quantification of HPL staining levels of Hela-IS cells co-expressing with wild type or mutant GBF1 without (blue bars) or with 4 hours imdz treatment (pink bars). Values were from three experimental replicates. Values on graphs indicate the mean ± SD. Statistical significance (p) measured by two-tailed paired t test.*, p < 0.05 and **, p<0.001 relative to untreated (0h) or GFP expressing cells. NS, non significant.

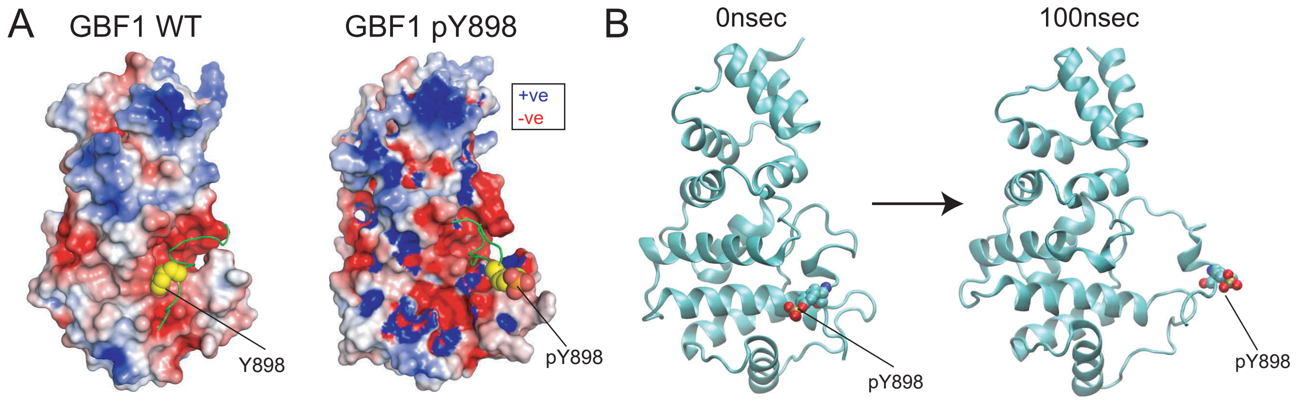


Supplementary Figure 6

(A) Electrostatic map of the charged residues on GBF1 Sec7d in the presence (right) and absence (left) of phosphorylation on Y898 on the C-terminal linker. (B) MD snapshot of the release of the C-terminal linker from the main body of the Sec7d when Y898 is phosphorylated. Values on graphs indicate the mean ± SD. Statistical significance (p) measured by two-tailed paired t test.*, $p < 0.05$ and **, $p < 0.001$ relative to untreated cells or to 10-min imdz treated cells expressing wild type GBF1. NS, non significant.

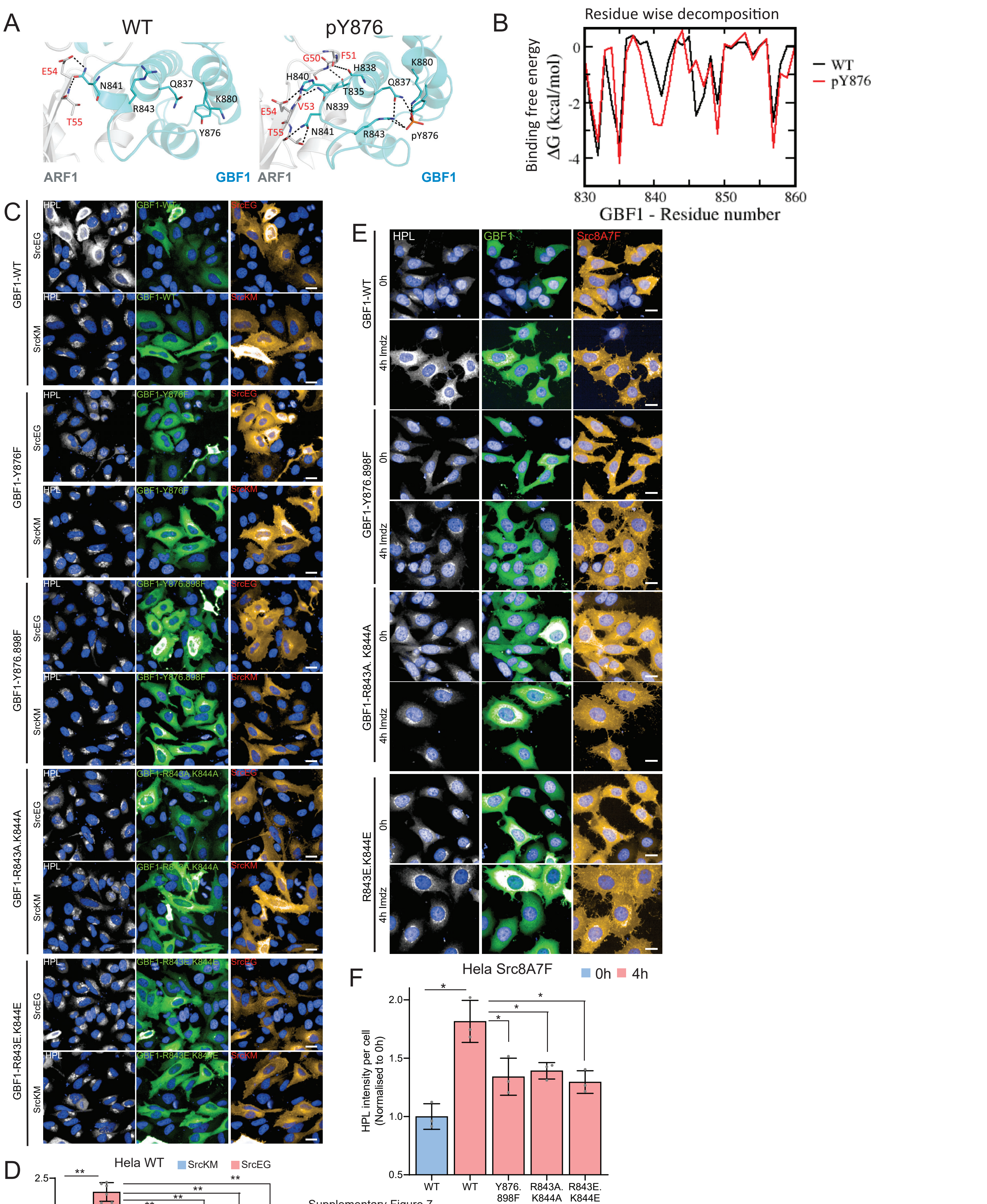

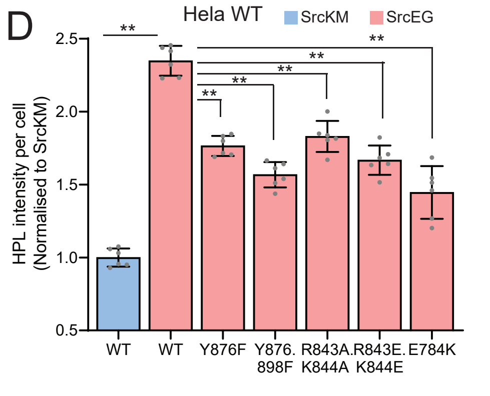


Supplementary Figure 7

(A) Representation of the predicted electrostatic bonds between the GBF1: Arf1 complex in the unphosphorylated (WT) and Y876 phosphorylated states. The Sec7d was shown in blue while Arf1 protein in grey. Residues involved in inter-protein interactions are shown as sticks, with inter-protein H-bonds highlighted as black dashed lines. (B) Per-residue decompositions of the binding free energies of interactions between the Y876-phosphorylated GBF1 Sec7d and Arf1 and between unphosphorylated GBF1 Sec7d and Arf1, using the MMGBSA approximations averaged over the conformations generated from MD simulations of the complexes. (C) Representative images of HPL staining in Hela cells co-expressing wild type or HI loop mutants with active SrcEG. Scale bar: 50 μ m. (D) Quantification of HPL staining levels of cells co-expressing wild type or mutant GBF1 with inactive SrcKM (blue bar) or active SrcEG (pink bars). Values were from six replicates from two independent experiments. (E) Representative images of HPL staining in Hela-IS cells co-expressing wild type or HI loop mutants. Scale bar: 50 μm. (F) Quantification of HPL staining levels of Hela-IS cells co-expressing wild type or mutant GBF1 that were unstimulated (0h, blue bar) or stimulated with 5 mM imdz (4h, pink bars). Values were from three replicates. Values on graphs indicate the mean ± SEM. Statistical significance (p) measured by two-tailed paired t test.*, p < 0.05 and **, p<0.001 relative to control cells. NS, non significant.
